# Allosteric pathways govern Gα protein coupling selectivity at promiscuous GPCRs

**DOI:** 10.64898/2026.08.28.747882

**Authors:** Tegvir S. Boora, Hanyu Chen, Aaron Cho, Wijnand J. C. van der Velden, Yoon Namkung, Yubo Cao, Jay Y. Paranjpe, Franziska M. Heydenreich, Andrei S. Rodin, Sergio Branciamore, Nagarajan Vaidehi, Stéphane A. Laporte

## Abstract

G protein-coupled receptors (GPCRs) regulate diverse physiological responses by engaging distinct heterotrimeric G proteins, yet the basis of Gα selectivity in promiscuous receptors remains unclear. Although ligand bias holds therapeutic promise, selectivity has been assumed to reside mainly in the ligand-binding site (LBS) or G protein interface (GPI). Here, we combine whole-receptor mutagenesis, functional Gα protein assays, molecular dynamics simulations, interpretable machine learning method, and Bayesian network modeling to identify residue networks governing Gα_q/11_ and Gα_12/13_ coupling and to define the molecular basis of G protein preference and promiscuity at two vasopressor GPCRs, the angiotensin II type 1 and prostaglandin F2α receptors. While residues within the LBS and GPI domains contribute to coupling efficiency and subtype discrimination, we find that long-range allosteric communication across the receptor, including from structurally unresolved domains, is the principal determinant of Gα protein preference and promiscuity. These allosteric pathways integrate multiple receptor domains, confer signaling robustness to mutation, and hierarchically govern coupling preferences. Our findings suggest that Gα protein selectivity is an allosterically encoded property of GPCRs and provide a conceptual framework for designing ligands and receptors with tailored Gα protein-biased signaling.

## INTRODUCTION

G protein-coupled receptors (GPCRs) are the largest family of membrane receptors and play a central role in transducing extracellular signals into intracellular responses^1–3^. GPCRs regulate a broad spectrum of physiological functions and constitute the largest class of therapeutic targets, with more than one-third of approved drugs acting on this receptor family^4^. They signal through interactions with heterotrimeric G proteins consisting of Gα and the Gβγ protein dimer^1–3^. Gα consists of four major subfamilies, Gα_q/11_, Gα_12/13_, Gα_s/olf_, and Gα_i/o_, with many receptors capable of engaging more than one Gα protein subtype, thereby triggering different signaling cascades and hence distinct physiological effects^5–7^. In addition, most GPCRs engage β-arrestins, which modulate receptor desensitization, internalization, trafficking, and signaling^8^.

GPCRs exhibit selective coupling to the Gα protein subtypes or β-arrestin whereby a ligand-GPCR pairing preferentially activates one signaling pathway over another^9,10^. This phenomenon of biased signaling or functional selectivity has been leveraged to develop signaling pathway-specific therapeutics with enhanced efficacy. While most approaches have focused on biasing GPCRs between Gα protein and β-arrestin pathways, recent interest has arisen in exploiting functional selectivity among different Gα proteins at promiscuous GPCRs, opening avenues for therapeutic applications^11–17^. Residues within the receptor-G protein-interacting interface (GPI) and long-range allosteric communication between the ligand-binding site (LBS), extracellular (EC), and intracellular (IC) regions are known to contribute to Gα protein-coupling^5,18–20^. However, how domains and residues in receptors encode specificity for distinct Gα subtypes (e.g. Gα_s/olf_, Gα_q/11_, Gα_i/o_, Gα_12/13_) remains unresolved, therefore limiting the rational design of functionally selective GPCR ligands.

GPCRs are inherently dynamic, sampling multiple conformational states that enable G protein binding^2,3,21^. Ligands, including biased ones, are thought to stabilize specific receptor conformations, thereby promoting preferential coupling to distinct Gα proteins. Structural methods such as X-ray crystallography and cryo-EM have provided valuable snapshots of ligand- and Gα protein-bound GPCR conformations, but they are limited in capturing the full dynamic complexity of these receptors^13,22–25^. Molecular dynamics (MD) simulations have revealed that multiple receptor regions, including the LBS, EC loops (ECLs), transmembrane (TM) helices, IC loops (ICLs), as well as the GPI, cooperatively and differentially contribute to Gα protein binding and activation through allosteric communication to one another^5,13,20,26,27^. Yet, the actionable information that can be extracted from these allosteric networks in a data-driven fashion to (i) uncover the differences in the allosteric communication mechanisms in the active versus inactive states of the receptor and (ii) provide a systematic method that characterizes the role of each amino acid in the GPCR in regulating its coupling to different Gα proteins is lacking. Additionally, how allostery shapes the shared and distinct coupling preferences of two promiscuous GPCRs engaging two Gα proteins remains unknown.

In this work, to uncover such allosteric communication mechanisms governing Gα protein preference at promiscuous GPCRs, we used two vasopressor receptors: the angiotensin II type 1 receptor (AT1R) and the prostaglandin F2α receptor (FP). Both couple to Gα_q/11_ and Gα_12/13_ with different coupling strengths, mediating small G protein Rho activation and driving smooth muscle contraction and regulating vascular tone^6,28–33^. While Gα_q/11_ interactions with these receptors have been structurally resolved, no such structural data exist for Gα_12/13_ binding with these receptors^34,35^.

We performed systematic alanine scanning of both receptors and assessed Gα_q/11_- and Gα_12/13_-mediated signaling via shared downstream effectors. This unbiased approach identified residues and domains including the disordered regions of receptors that regulate G protein preference, promiscuity, and resilience to mutations. Mechanistic insights into the functional data for both receptors were further interpreted by integrating molecular dynamics (MD) simulations and an interpretable machine learning model, Bayesian network modeling (BNM). In contrast to correlation-based dynamic network analyses, which detect co-fluctuations but lack causal directionality, BNM infers conditional dependencies among residues^36,37^. This approach enabled us to identify directed allosteric couplings and communication pathways linking each receptor’s LBS to their GPI^5,13,22,26,27^. This combined analysis provides a holistic approach to uncover the role of the dynamic and disordered elements of the receptor in regulating the differential coupling of the two receptors to two different Gα proteins. In an accompanying study, we used these approaches and identified a negative allosteric modulator to AngII-mediated activity of AT1R^38^. Here, by integrating computational modeling with mutagenesis, we show that Gα protein preference and resilience extend beyond direct receptor-G protein contacts and are shaped by long-range allosteric communication and cooperative residue networks spanning the LBS and structurally unresolved receptor domains. These findings reveal mechanisms of Gα protein selectivity in shared effector pathways, define a hierarchy of signal efficiency at promiscuous receptors, and provide a framework for rational design of biased receptors or ligands.

## RESULTS

### Alanine scanning mutagenesis of FP and AT1R reveals domain-specific control of cell surface expression and differential Gα_q/11_ vs. Gα_12/13_ coupling sensitivity

Expanding on our prior alanine-scanning mutagenesis of AT1R^26^, we substituted each FP receptor residue with alanine or glycine in the case of native alanine, creating a new library of 358 single-point mutants to examine the determinants of Gα_q/11_ and Gα_12/13_ protein selectivity (see Methods). We validated cell-surface expression of the FP mutants, as previously done for the AT1R mutant library^26^, and considered mutants inadequately expressed for functional analysis if expression was below 34% and 50% of WT levels for FP and AT1R, respectively, and was accompanied by significant reductions in Gα_q/11_ and Gα_12/13_ signaling (Supplementary Fig. 1a, b). Of 358 mutants per receptor, 19 FP (5%) and 22 AT1R (6%) with sub-threshold expression were excluded, leaving 339 FP and 336 AT1R mutants for functional analysis (Supplementary Fig. 2a, b). Many point mutations that increased cell surface expression receptor to nearly or more than two-fold that of WT were identified at each receptor (FP: L9A^N-term^, S118A^3.35^, T268A^6.54^, L17A^N-term^, L228A^5.64^, etc.; AT1R: T216A^5.59^, F77A^2.53^, S347A^C-tail^, V116A^3.40^, R137A^ICL2^, etc.; Ballesteros-Weinstein (BW) numbering or receptor domain in upper case)^39^ (Supplementary Fig 2a, b). For FP, expression-potentiating mutations were localized in structurally unresolved regions like the N-terminus of the receptor, while for AT1R, they were widely distributed in the transmembrane (TMs), and the intracellular face of the receptor (e.g.; ICL2, ICL3, and helix-8 (H8)) and the C-terminus) (Supplementary Fig 2a), suggesting a receptor- and region-specific regulation of cell surface expression.

We next profiled the pharmacological properties of mutant FP and AT1R receptors with respect to Gα protein coupling preference using their respective endogenous ligands, PGF2α and AngII. Building on our previous AT1R study, in which Gα_q/11_ signaling was measured using a BRET-based PKC biosensor in parental HEK293 cells, we assessed FP mutant signaling using a BRET-based Rho effector biosensor in cells lacking (Knock-Out, KO) either Gα_q/11_ or Gα_12/13_. Gα_12/13_ coupling of AT1R mutants was similarly measured with the Rho biosensor in the presence of the G_αq/11_ inhibitor YM-254890 within parental cells. These approaches allowed us to isolate the individual Gα_q/11_ and Gα_12/13_ pathways, as Rho signaling can be activated downstream of both G protein families^26,29,40,41^. Pertussis toxin (PTX) treatment confirmed that Gα_i/o_ does not contribute to FP receptor- or AT1R-mediated Rho signaling in our paradigm (Supplementary Fig. 3)^29^. Concentration-response curve analyses yielded maximal response (Emax), half-maximal effective concentration (EC_50_), and activity (log(Emax/EC_50_)) of each mutant for each pathway compared to WT receptors (Supplementary Data 1, 2). Downstream activation of a subset of FP and AT1R mutants shows strong correlation between mutation-dependent effects and Gα protein coupling, using Gα_q_ or Gα_13_ Gαβγ (GABY) BRET sensors as well as reproducibility of mutagenic signaling effects in different biosensor paradigms, using the Rho biosensor in Gα_q/11_ and Gα_12/13_ KO cells (Supplementary Fig. 4a, b; Supplementary Data 3, 4)^26^. Consistent effects across these assays indicate that the downstream effector BRET-based biosensor response accurately reports GPCR-Gα_q_/Gα_13_-protein activation of mutants. Mutants were therefore classified as unaffected (not significantly different from WT) or affected (gain or loss of function relative to WT) for the Gα_q/11,_ Gα_12/13_ or both pathways based on the pharmacological parameters of Emax, EC_50_, activity, or combinations thereof (Fig. 1a). Distinct receptor-specific patterns of Gα protein sensitivity to point mutations emerged between FP and AT1R (Fig. 1a). For FP, the Gα_q/11_ signaling pathway was more sensitive to point mutations than Gα_12/13_, whereas AT1R showed the opposite trend. FP-Gα_q/11_ and AT1R-Gα_12/13_ displayed comparable mutational sensitivities with 53% and 56% of mutants functionally affected, while FP-Gα_12/13_ and AT1R-Gα_q/11_ pathways were more mutation-resilient with only 33% and 27% of receptor mutants affecting responses. These reciprocal patterns reveal receptor-specific, pathway-specific determinants of Gα protein signaling, which was evident from the Emax and EC_50_ sensitivities of mutant receptors at the Gα_q/11_ or Gα_12/13_ pathways (Fig 1a).

**Figure 1.**
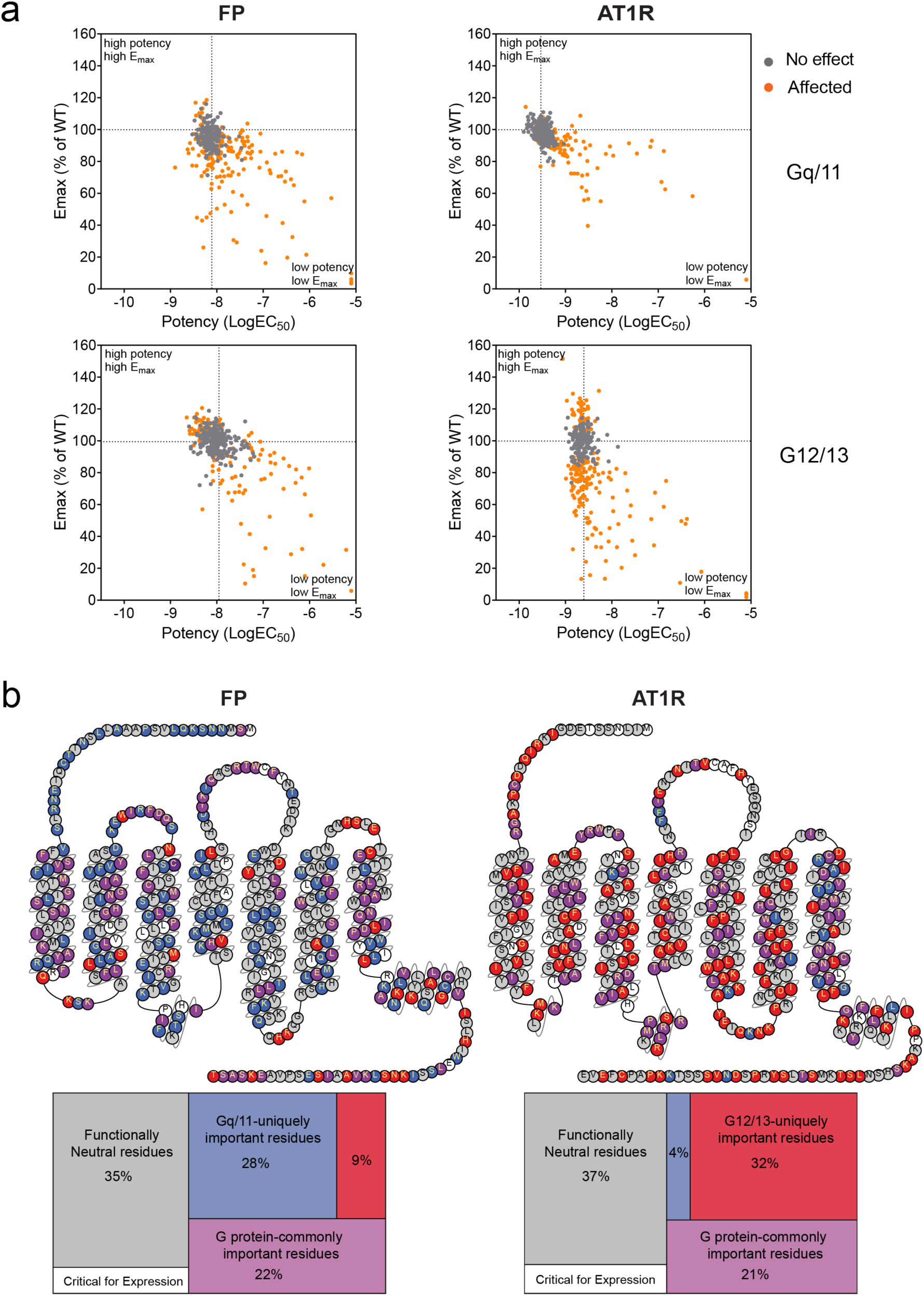
Mutational landscape of FP and AT1 receptors reveals Gα protein signaling preference. **a** Effects of alanine/glycine substitutions on FP (left) and AT1R (right) signaling through Gα_q/11_ (top) and Gα_12/13_ (bottom). Scatter plots show signaling properties of individual FP and AT1R mutants relative to their corresponding wild-type (WT) receptors. Mutants are categorized as unaffected (grey) or functionally affected (orange) relative to WT. Insets indicate the fraction of WT-like versus affected variants. A mutation was classified as affected if any pharmacological parameter (Emax, logEC_50_, and/or relative activity [RA]) differed significantly from WT (*q* < 0.05). Grey dotted lines denote WT reference values for efficacy and potency. Data represent means from ≥3 independent experiments; pharmacological parameters were derived from full concentration-response curves. **b** Functional categorization and topological mapping of residues contributing to Gα_q/11_ and Gα_12/13_ coupling in FP (left) and AT1R (right). Residues classified as Critical for Expression are shown in white lettering, reflecting reduced cell-surface expression and signaling decreases in both pathways upon alanine/glycine substitution. Functionally neutral positions (grey) exhibited no detectable effect on signaling upon mutation. G protein-commonly important residues, whose mutation significantly affected both Gα_q/11_ and Gα_12/13_ signaling, are shown in purple, whereas Gα_q/11_-uniquely important Gα_12/13_-uniquely important residues, whose mutation significantly affected one pathway but not the other, are shown in blue and red, respectively. Snake plots were adapted from GPCRdb (www.gpcrdb.org).

We next categorized residues based on these aforementioned functional effects of point mutations at individual sites, taking into consideration both pathways, and established five categories of residues: (1) Critical for Expression; (2) Functionally Neutral; (3) Gα protein-commonly important; (4) Gα_q/11_-uniquely important; (5) Gα_12/13_-uniquely important and mapped them onto their respective receptors (Fig. 1b). Gα_q/11_-uniquely important and Gα_12/13_-uniquely important refer to residues whose mutation significantly affected signaling through one G protein pathway but not the other under our predefined criteria. These classifications describe pathway-specific mutational effects and do not represent measures of signaling bias. Categories 3, 4, and 5 comprised the ensemble of Functionally Important residues that caused significant changes in mutant signaling relative to WT. Mutations of the residues that are Critical for Expression severely impaired cell surface expression, indicating potential roles in folding and trafficking, while all other mutants were well expressed at the cell surface. Functionally Neutral residues had no significant effect on Gα_q/11_ or Gα_12/13_ signaling, as compared to WT. Gα protein-commonly important residues altered responses in both pathways upon mutation, therefore inferred to be functionally important for coupling to both Gα protein families. Interestingly, similar percentages and distributions of these three residue classes were observed in both receptors, suggesting evolutionarily conserved proportions of residues, whose roles in expression, signaling, and functional neutrality are conserved across Class A GPCRs. However, we identified functionally important residues whose mutation uniquely affected either Gα_q/11_ or Gα_12/13_ signaling. These Gα_q/11_ or Gα_12/13_-uniquely important residues may help define the molecular determinants of G protein coupling selectivity in promiscuous GPCRs. Interestingly, their relative number, localization, and effects for each pathway at each receptor differed markedly. In FP, Gα_q/11_-uniquely important residues (28%) were spread throughout the receptor, while Gα_12/13_-uniquely important residues (9%) were fewer in number than for Gα_q/11_ and mainly clustered in TM7, H8, and the C-terminus. AT1R showed the reverse trend, with few Gα_q/11_-uniquely important (4%) mainly clustered in TM7 but many Gα_12/13_-uniquely important residues (32%) which were distributed throughout the receptor. In both receptors, mutation of predominantly Gα_12/13_-uniquely important C-terminal residues enhanced Gα_12/13_ signaling, revealing a conserved inhibitory role for this structurally unresolved region in Gα_12/13_ coupling. These findings suggest that Gα signaling is governed by a conserved activation core supporting common coupling and pathway-uniquely important residues that may drive Gα protein selectivity. Notably, about two-thirds of residues at FP and AT1R were found to be functionally important, many of which localized to disordered or unresolved receptor regions, revealing previously underappreciated functional determinants only identifiable through systematic mutagenesis.

### LBS and GPI residue interactions in FP and AT1R partially determine Gα protein coupling and preference

As the LBS and GPI are central to Gα protein signaling^1,7,19,42,43^, with the GPI contributing to Gα coupling selectivity as shown by us and others^18,19^, we next assessed how residues within these domains contribute to functional selectivity. Although structures are available for each FP and AT1R bound to their endogenous ligands and WT Gα_q_, no structures exist with any Gα_13_ isoform in complex with either receptor. Importantly, these structures represent static conformations and capture only a subset of the most stable ligand-receptor and receptor-Gα protein contacts. Given that ligand-receptor and receptor-Gα protein interactions are highly dynamic, we reasoned that a state-resolved, residue-level analysis was required to better define the determinants of Gα coupling selectivity. We thus performed all-atom MD simulations on the FP and AT1R bound to their endogenous ligands (PGF2α and AngII, respectively) and either Gα_q_βγ or Gα_13_βγ in a POPC lipid bilayer (see Methods for details). For each conformational state, we performed five independent 1 μs MD simulations and uncovered the convergence (Supplementary Fig. 5). We calculated the frequencies of the contacts between the receptor-ligand and receptor-Gα protein along the trajectories. The near totality of high frequency contacts observed in our simulations were also found to be present in the cryo-EM structure interactions^34,35^, while lower-frequency contacts revealed additional, previously unresolved contact interfaces between each receptor and their respective ligand, as well as Gα_q_/Gα_13_ (Supplementary Fig. 6, 7), unobserved in structures.

We defined the LBS as residues that formed MD-revealed contacts with the endogenous ligand with a contact-frequency threshold of 0.2: PGF2α for FP and AngII for AT1R (see Methods for details), to minimize false positives (Supplementary Fig 6, 7). We compared ligand-residue contacts between Gα_q_- and Gα_13_-coupled states in FP and AT1R, respectively, to identify shared vs. unique residue interactions in each receptor. Within the LBS, 24 residues showed contact with PGF2α in the FP receptor and 34 residues with AngII in AT1R, with most of these ligand-residue contacts shared across Gα_q_ and Gα_13_ complexes of the same receptor (Fig. 2a; Supplementary Data 5a, b). However, a subset of LBS residues formed ligand contacts exclusively in either the Gα_q_- or Gα_13_-bound state (6 Gα_q_-specific and no Gα_13_-specific contacts for FP; 4 Gα_q_-specific and 2 Gα_13_-specific contacts for AT1R), namely FP-PGF2α Gα_q_-specific: S33^1.39^, F36^1.41^, M37^1.43^, F111^3.28^, A293^7.42^, Q297^7.46^; and AT1R-AngII Gα_q_-specific: F77^2.53^, W253^6.48^, A291^7.42^, Y292^7.43^; and AT1R-AngII Gα_13_-specific: S105^3.29^, S109^3.33^ (Fig. 2a; Supplementary Data 5a, b). These state-specific contacts suggest that the LBS contributes to G protein functional selectivity and may be exploited to design functionally selective ligands. Within the LBS, the conserved tryptophan of the CWxP motif was identified as a ligand-contacting residue in both FP (W262^6.48^) and AT1R (W253^6.48^). In FP, W262^6.48^ additionally forms part of the LLW motif^34^, a known hydrophobic core triad within FP, further underscoring the conserved importance of this position in ligand-GPCR interactions.

**Figure 2.**
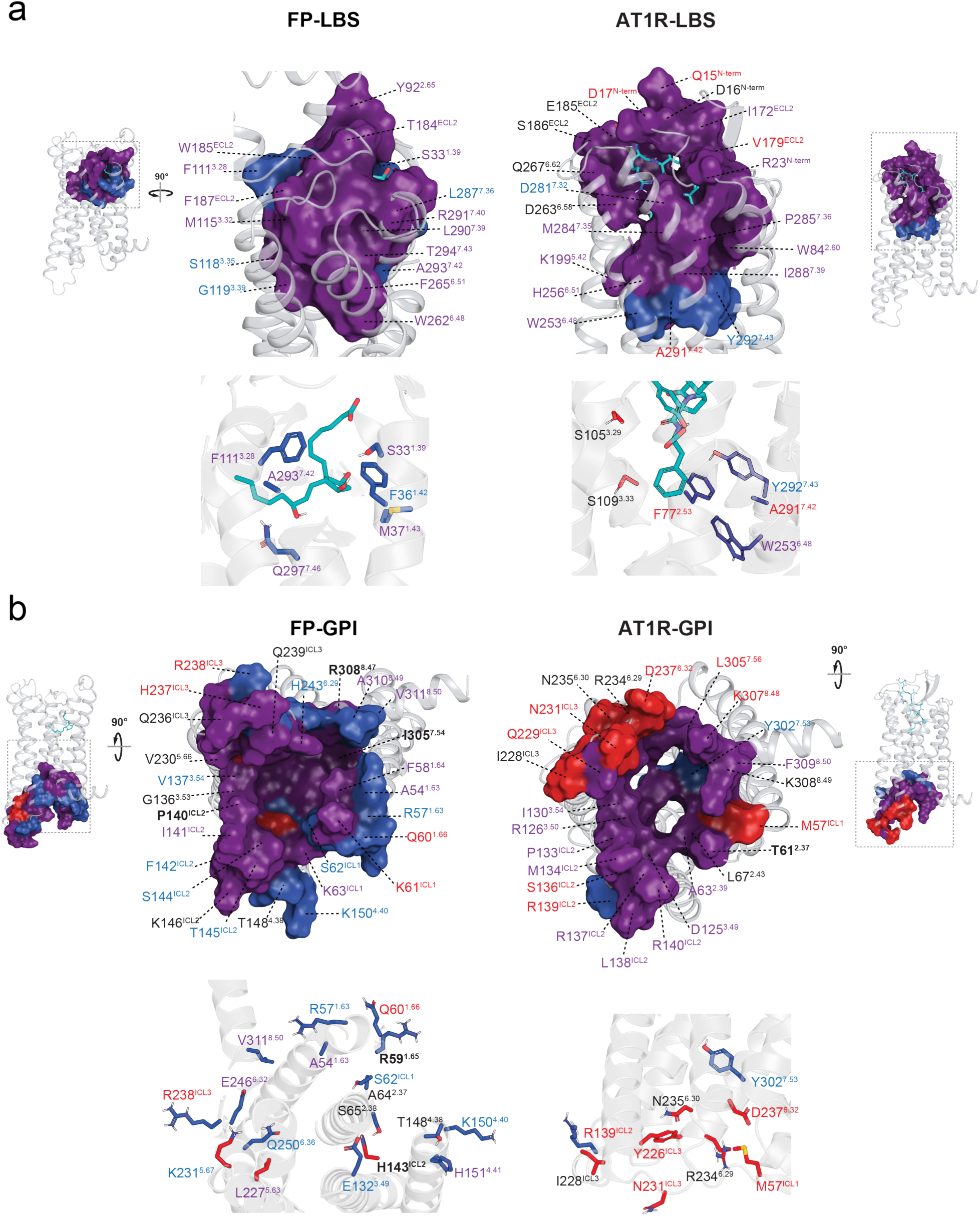
Structural mapping of ligand-binding sites (LBS) and G protein coupling interfaces (GPI) in FP and AT1R reveals common and Gα protein-specific coupling determinants. **a** Residues in the ligand binding sites showing contacts with PGF2α in FP and AngII in AT1R calculated from MD simulations of PGF2α-bound FP (left) and AngII-bound AT1R (right), each in complex with either Gα_q_ or Gα_13_. Residues in the LBS that contact the endogenous ligand in both Gα_q_- and Gα_13_-bound conformations are shown in purple, whereas residues that contact the ligand only in the Gα_q_ -bound state are shown in blue (top). The contact points of these Gα protein-bound-state-specific ligand-residue contacts in the LBS with their respective ligands for each receptor are shown in ribbon rendering with their residue numbers in the GPCR residue numbering system (bottom). **b** Residues in the Gα protein interface (GPI) that form contacts with Gα_q_, Gα_13_, or both in FP (left) and AT1R (right) calculated from MD simulations of PGF2α-bound FP and AngII-bound AT1R, each in complex with either Gα_q_ or Gα_13_. Residues contacting both Gα_q_ and Gα_13_ are shown in purple (top), whereas residues exhibiting Gα protein-specific contacts are shown in blue (Gα_q_) and red (Gα_13_). Gα protein-specific residue contacts are additionally displayed in ribbon representation, with positions labeled using the GPCR generic residue numbering scheme (bottom). In the LBS and GPI, residue labels shown in purple indicate functionally Gα protein-commonly important residues, those in blue indicate Gα_q_-uniquely important residues, and those in red indicate Gα_13_-uniquely important residues. Bolded labels indicate residues classified as Critical for Expression.

Likewise, the GPI was defined as residues that established persistent contacts with one or both Gα proteins (Gα_q_ and/or Gα_13_), above the established 0.2 frequency cutoff, in MD simulations as we have done previously^5^ (see Methods). Within the GPI, 47 residues of FP and 33 residues of AT1R showed contact with either or both Gα_q_ and Gα_13_ (Fig. 2b; Supplementary Data 5c, d). A majority of Gα protein-residue contacts were shared between both Gα_q_ and Gα_13_, and an important minority were specific to either Gα protein (17 Gα_q_-specific and 3 Gα_13_-specific for FP and 3 Gα_q_-specific and 8 Gα_13_-specific for AT1R), namely for FP-Gα_q_ specific: A54^1.60^, R57^1.63^, R59^1.65^, Q60^1.66^, S62^ICL1^, A64^2.37^, S65^2.38^, E132^3.49^, T148^4.38^, K150^4.40^, H151^4.41^, R238^ICL3^, G240^ICL3^, E246^6.32^, Q250^6.36^, A310^8.49^, V311^8.50^; for FP-G13 specific: H143^34.53^, L227^5.63^, K231^5.67^; for AT1R-Gα_q_ specific: R139^ICL2^, Y302^7.53^, G303^7.54^; for AT1R-G13 specific: M57^ICL1^, A225^5.68^, Y226^ICL3^, 228^ICL3^, N231^ICL3^, R234^6.29^, N235^6.30^, D237^6.32^) (Fig. 2b; Supplementary Data 5c, d), consistent with Gα protein selectivity mechanism regulation at each GPI^5^. Within the GPIs, residues from the DRY/ERC and NPxxY/DPxxY motifs were identified in both receptors: FP (R133^3.50^ in the ERC motif; Y304^7.53^ in the DPxxY motif) and AT1R (R126^3.50^ in the DRY motif; Y302^7.53^ in the NPxxY motif). These findings support a conserved role for these motifs in Gα protein coupling.

Across our analyses, an average of 93% (range, 87-100%) of ligand-residue and residue-G protein contacts identified in the FP and AT1R structures (8IUK and 7F6G)^34,35^ were recapitulated in our simulations, with the remaining contacts falling below the 0.2 contact-frequency threshold. Moreover, an average of 70% (range, 56-100%) of residues predicted to contact the ligand or G protein in our simulations exhibited altered signaling through the corresponding pathway and/or changes in receptor expression. These findings demonstrate that our MD simulations robustly capture experimentally defined receptor-ligand and receptor-G protein interfaces and identify contacts enriched in functionally important residues. Overall, these results support a model in which the LBS and GPI largely mediate the common coupling of a promiscuous GPCR to different Gα proteins, while also harboring discrete determinants that contribute to coupling promiscuity and preference, particularly within the GPI.

### Region- and residue-specific localization of Gα protein coupling determinants differs between two promiscuous GPCRs

To further parse the structural regions of FP and AT1R containing selectivity determinants, we mapped all residues that showed Gα protein-unique functional importance onto both FP and AT1R receptors. This analysis revealed that all TM helices relatively contribute comparably to selectivity at both receptors (Fig. 3a), except for AT1R’s TM6 having relatively more G protein-uniquely important residues. The core of the FP receptor predominantly contained Gα_q/11_-uniquely important residues while the core of AT1R was dominated by Gα_12/13_-uniquely important sites. The ECLs played a relatively larger role in Gα protein selectivity at FP than AT1R, with FP ECL1 primarily and ECL2 exclusively containing Gα_q/11_-uniquely important residues while ECL3 exclusively contained Gα_12/13_-uniquely important sites. In the case of AT1R, its ECL2 primarily contained Gα_12/13_-uniquely important residues, with no Gα protein-uniquely important residues found in its ECL1 or ECL3. FP contained a balanced proportion of Gα_q/11_-uniquely important vs Gα_12/13_-uniquely important residues across the ICLs (both in ICL1, only Gα_q/11_-uniquely important in ICL2, and only Gα_12/13_-uniquely important in ICL3) while AT1R’s ICLs were found to be near exclusively enriched in Gα_12/13_-uniquely important residues, demonstrating the importance of these GPCR unresolved domains in regulating Gα coupling promiscuity and selectivity. The N-terminus was found to impart opposing selectivity, containing Gα_q/11_-uniquely important residues in FP and Gα_12/13_-uniquely important residues in AT1R. The only shared feature between the two receptors was the role of the C-tail in suppressing Gα_12/13_ coupling. In both FP and AT1R, point mutations that enhanced Gα_12/13_ signaling clustered within the C-terminus, underscoring the importance of this unresolved region in restraining Gα_12/13_ activation in the WT receptor. Our analysis underscores the importance of residues that are both region-specific and receptor-specific in regulating Gα protein selectivity across promiscuous receptors, demonstrating no universal motifs governing Gα protein selectivity other than the common inhibitory role of each GPCR’s C-tail for Gα_12/13_ coupling.

**Figure 3.**
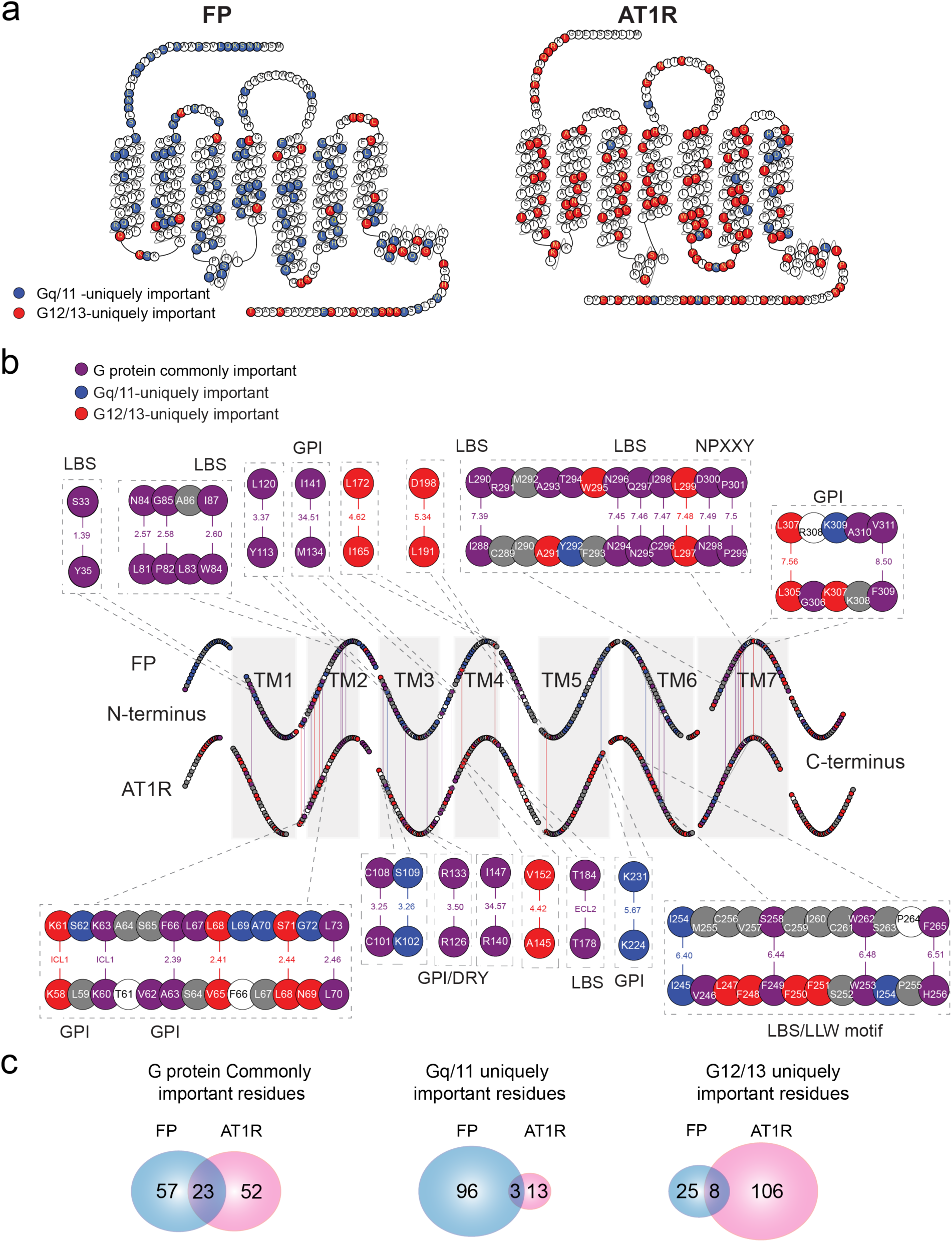
Gα protein-specific coupling determinants differ between receptors, while canonical structural domains support common coupling. **a** Location of Gα protein-uniquely important residues in FP (top) and AT1R (bottom) receptors. FP was primarily enriched in Gα_q/11_-uniquely important (blue) residues whereas AT1R was enriched in Gα_12/13_-uniquely important (red) residues. Snake plots are adapted from GPCRdb (www.gpcrdb.org). **b** Alignment of functionally important residues located at homologous positions in FP (top) and AT1R (bottom). Snake plots display all 358 residues (excluding the start and stop codons), arranged sequentially and colour-coded according to panel **c** of Figure 1. Homologous residues were aligned based on Ballesteros-Weinstein (BW) numbering and alignment of intracellular and extracellular loops; shared functional roles are connected by lines. Functionally homologous residues within conserved functional motifs or the LBS or GPI of both receptors are annotated. **c** Numbers of functionally homologous residues among functionally important residues in FP and AT1R. The Venn diagrams display the numbers of Gα protein-commonly important, Gα_q/11_-uniquely important, and Gα_12/13_-uniquely important residues in FP (blue) or AT1R (red). Purple numbers indicate homologous residues shared between FP and AT1R within each category.

We narrowed our focus to examine any possible functional homology at a residue-level between FP and AT1R. We defined functionally homologous residues as those in the same structural position in GPCRs according to the BW residue numbering system, which identifies conserved residue positions in resolved regions of GPCRs, and had the same functionally important role (Gα protein-commonly important, Gα_q/11_-uniquely important, or Gα_12/13_-uniquely important), as revealed by our alanine scanning approach. Alignment of all functionally important residues in resolved regions of FP and AT1R identified 34 functionally homologous residue positions. These residues are found within all TMs and H8, but primarily clustered in TM2 and TM7 s 1-3, 6-7, ICLs 1-2, ECL2, and helix 8 (Fig. 3b). Most functionally homologous residues resided within the LBS or GPI and were residues commonly functionally important to both Gα_q/11_ and Gα_12/13_. Functionally homologous residues that were uniquely functionally important for one Gα protein pathway are fewer: only 3 Gα_q/11_- and 8 Gα_12/13_-uniquely important residues were homologous, while most Gα protein uniquely important sites were non-homologous receptor-specific positions. Thus, while certain LBS- and GPI-based residues form a structurally similar core that enables general Gα protein coupling across both receptors, few functionally homologous Gα protein-uniquely important residues between both receptors were observed, therefore suggesting the determinants of Gα protein functional selectivity to be specific to different promiscuous GPCRs.

### Allosteric networks identified using Bayesian Network Model that are outside the LBS and GPI govern Gα Protein coupling preference

As residues acting as selectivity determinants represented only a minority within the LBS and GPI, each of which contained a conserved core mediating shared Gα coupling across both receptors, we hypothesized that Gα preference is primarily encoded by distributed allosteric networks that integrate long-range intramolecular communication throughout the receptor. Consistent with this hypothesis, approximately three-quarters of functionally important residues in both receptors were located outside the LBS and GPI and were therefore considered putatively allosteric (Fig. 4a).

**Figure 4.**
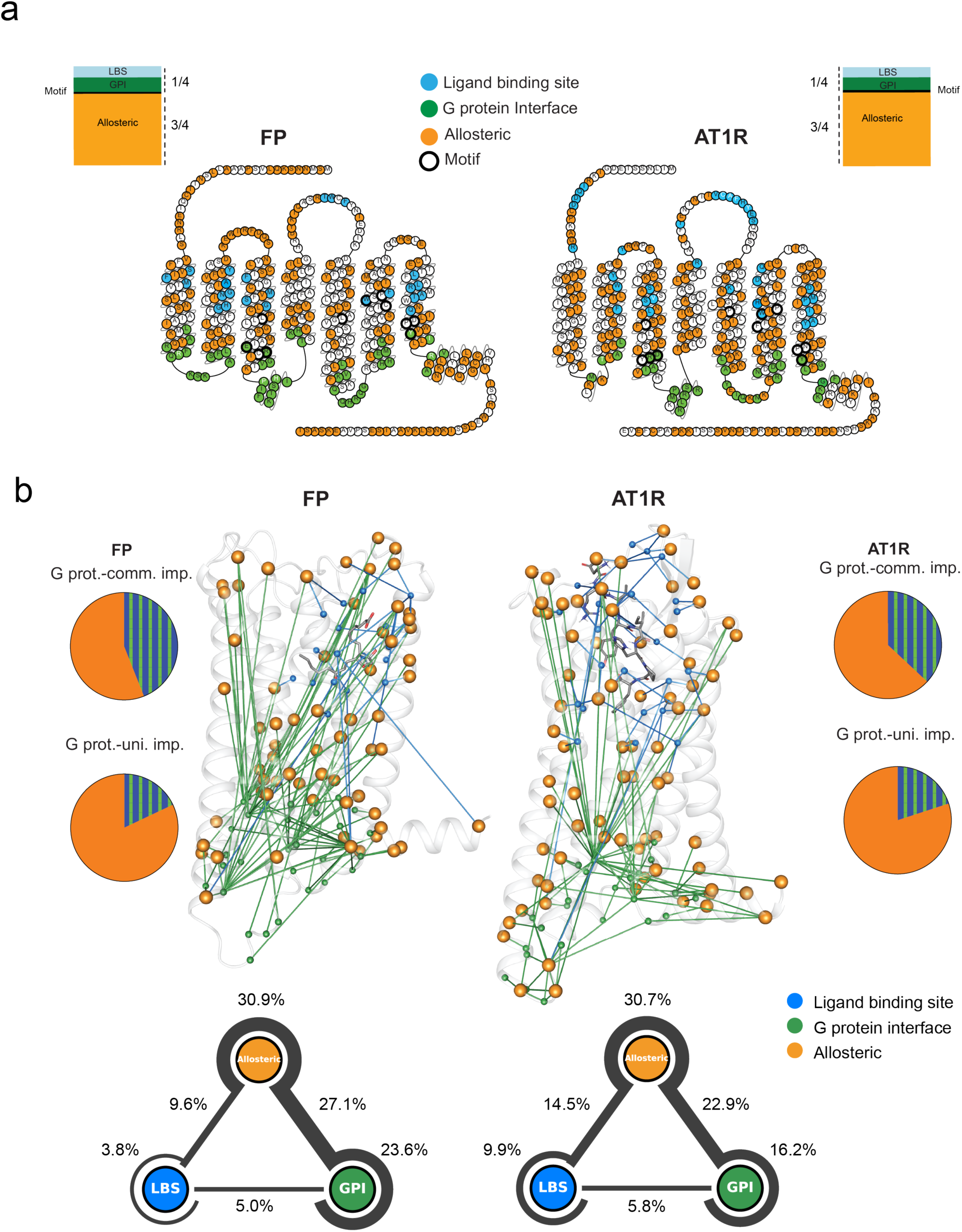
Allosteric networks are major determinants of Gα protein coupling selectivity in FP and AT1R. **a** Topological distribution of functionally important residues (Gα protein-commonly important, Gα_q/11_-uniquely important, Gα_12/13_-uniquely important) in FP (left) and AT1R (right) from the BRET measurements of Gα_q/11_ and Gα_12/13_ coupling. Functionally important residues within the Ligand Binding Site (LBS) (blue), G protein-binding interface (GPI) (green), within known functional motifs (e.g. DRY & NPxxY motifs) (bolded) or inferred to be allosteric (orange) are mapped onto snake plots (top) of FP and AT1R. Residues within the LBS and GPI critical for expression are written in white. Proportions of functionally important residues localized within the LBS, GPI, known structural functional motifs, or outside these defined domains (bottom) show that approximately one-quarter of functionally important residues localize to the LBS/GPI or known functional motifs, whereas approximately three-quarters lie outside these regions. Snake plots are adapted from GPCRdb (www.gpcrdb.org). **b** Molecular dynamics (MD) simulations of functionally important residues forming allosteric connection pathways between interdomains of FP (top left) and AT1R (top right). Top-ranked functionally important residues in MD simulations having been inferred to be allosteric in functional mechanism in panel **a** are mapped onto the structure of FP (left) and AT1R (right) having been identified to allosterically (orange) connect the LBS and GPI. The ensemble of residue-to-residue connections is quantitatively represented in triangles for FP (middle left) and AT1R (middle right) with percentages of connections directly connecting LBS (blue) and GPI (green), bridging connection with the allosteric network (orange), or connections within these domains. The proportion of Gα protein-commonly important residues (top) and -uniquely important residues (bottom) which are either allosteric in mechanism or within LBS/GPI is represented as pie charts for FP (left) and AT1R (right).

We previously showed that many of these AT1R residues, extending beyond conserved functional motifs such as DRY, PIF, and NPxxY, form extensive networks connecting structurally distant receptor regions during Gα_q_ coupling^44^. These results thus support long-range allosteric communication as a determinant of Gα functional selectivity, with residues within these networks contributing uniquely or commonly to Gα_q/11_ and Gα_12/13_ coupling.

To concretely identify and elucidate these putative allosteric communication pathways for either or both Gα_q_ and Gα_13_ at each receptor, we used an interpretable and unsupervised machine learning model, the Bayesian network models (BNMs), to analyze the MD simulation trajectories using our software BaNDyT^45^. We previously used BNMs to predict and prospectively test the amino acid residue contacts that show high cooperativity and their allosteric effect in the interface of GPCR:G protein complexes^20,46,47^. Here, we used the nonbonded interaction energy between each residue and its surrounding residues within 12Å radius in the receptor structure, calculated for every MD snapshot, as input variables to generate the BNM for FP and AT1R. In the resulting residue-level BNMs analysis, which we have elaborated upon in a recent associated paper^38^, receptor residues are the nodes, the edges connecting two nodes show correlated movement or co-dependencies in residue dynamics, and edge weights are the strength of the co-dependencies.

In this work, we first generated two separate active-state BNMs using active-state MD trajectories of Gα_q_-bound and Gα_13_-bound complexes for FP and AT1R, respectively (Supplementary Data 6-9), and an inactive-state BNM from inactive-state MD simulations of FP and AT1R (Supplementary Data 10, 11) (see Methods for details). In each BNM, the weighted degree (Wd) of a residue is defined as the sum of the weights of all edges connected to that node. To validate the Gα_q_- and Gα_13_-BNMs, we tested whether state-dependent changes in Wd (ΔWd, active vs. inactive) could recover experimentally determined functional residues. All state-specific networks exhibited strong correspondence between ΔWd and alanine mutational effects on signaling output (EC_50_, Emax, or RA), with significant enrichment of ala-scanning positives (Supplementary Fig. 8). To assess co-dependencies present when either Gα protein was bound to the receptor, we constructed universal Gα_q_ - Gα_13_ BNMs for FP and AT1R (see Methods and Supplementary Data 12, 13). Co-dependencies involving functionally defined allosteric residues accounted for 67.6% and 68.1% of the total co-dependencies in FP and AT1R, respectively (Fig. 4b). Intra-allosteric co-dependencies constituted 30.9% in FP and 30.7% in AT1R, demonstrating extensive communication among functionally important residues outside the direct ligand- and G-protein-contact interfaces. By contrast, direct co-dependencies between LBS and GPI residues were uncommon, representing only 5.0% and 5.8% of the total co-dependencies in FP and AT1R respectively. Allosteric-LBS and allosteric-GPI co-dependencies together accounted for similar proportions in FP and AT1R (36.7% and 37.4%, respectively), consistent with potential indirect communication between the two interfaces through the surrounding allosteric network.

The distribution of these co-dependencies nevertheless differed between receptors. AT1R contained greater proportions of allosteric-LBS co-dependencies (14.5% vs. 9.6%) and intra-LBS co-dependencies (9.9% vs. 3.8%) than FP, indicating slightly stronger integration of the ligand-binding pocket with its surrounding receptor network. Conversely, FP contained greater proportions of allosteric-GPI co-dependencies (27.1% vs. 22.9%) and intra-GPI co-dependencies (23.6% vs. 16.2%), indicating a more GPI-centered network organization. Structural mapping of the co-dependencies between allosteric residues to both interfaces further showed that these relationships extend throughout the receptor. Together, these patterns are consistent with perturbations at the ligand-binding pocket or G-protein interface being coupled to a receptor-wide allosteric network rather than remaining confined to the respective contact site.

Cross-referencing the mechanistic location of each functionally important residue (LBS, GPI, or allosteric) with its experimentally determined role (G protein-commonly important, Gα_q/11_-uniquely important, or Gα_12/13_-uniquely important; Fig. 1) revealed distinct distributions between residues supporting common and Gα protein-specific coupling (Fig. 4b). Among G protein- commonly important residues 57% in FP and 66% in AT1R were allosteric, with the remaining residues located in the LBS or GPI. Allosteric residues were even more enriched among positions uniquely important for Gα_q/11_ or Gα_12/13_ coupling, accounting for 83.2% in FP and 82.2% in AT1R. Thus, residues associated with Gα protein selectivity are predominantly located outside the direct ligand- and G-protein-contact interfaces, supporting a major contribution of distributed allosteric networks to receptor function and Gα protein-selective coupling in both receptors.

### BNM connectivity within receptors uncovers Gα protein coupling preference and confers redundancy and resilience to mutations

We next posited that differences in the allosteric communication network of each GPCR between the Gα_q_-coupled vs. Gα_13_-coupled states to be the primary drivers of the GPCR promiscuity and selectivity. To investigate this, we compared Gα_q_-coupled vs. Gα_13_-coupled edge strengths in the “rescored” using mutual information (MI) in the universal BNM of each receptor (Supplementary Data 14-17). In the FP receptor, a larger proportion of edges was stronger in the Gα_13_-coupled state than in the Gα_q_-coupled state, whereas AT1R displayed the opposite pattern (Fig. 5a). In FP, 56% of edges exhibited stronger weights in the Gα_13_-bound state, whereas 44% were stronger in the Gα_q_-bound state. In contrast, AT1R showed the opposite distribution, with 59% of edges stronger in the Gα_q_-bound state and 41% stronger in the Gα_13_-bound state. Thus, each receptor displays a state-dependent redistribution of edge strengths within the allosteric network, whereby one Gα-bound state is characterized by a greater proportion of strong edges relative to the other. Notably, these trends paralleled experimental mutation resilience measured by BRET: FP-Gα_13_ and AT1R- Gα_q_ exhibited greater tolerance to point mutations than their respective counterparts. These results suggest that Gα protein-bound states enriched in system-prone edges form more densely interconnected allosteric networks, providing redundant communication routes between the LBS and GPI. Such redundancy confers resilience to single-residue perturbations. In contrast, sparser networks, such as FP-Gα_q_ and AT1R-Gα_13_, appear more vulnerable to disruption by individual mutations along the allosteric pathway.

**Figure 5.**
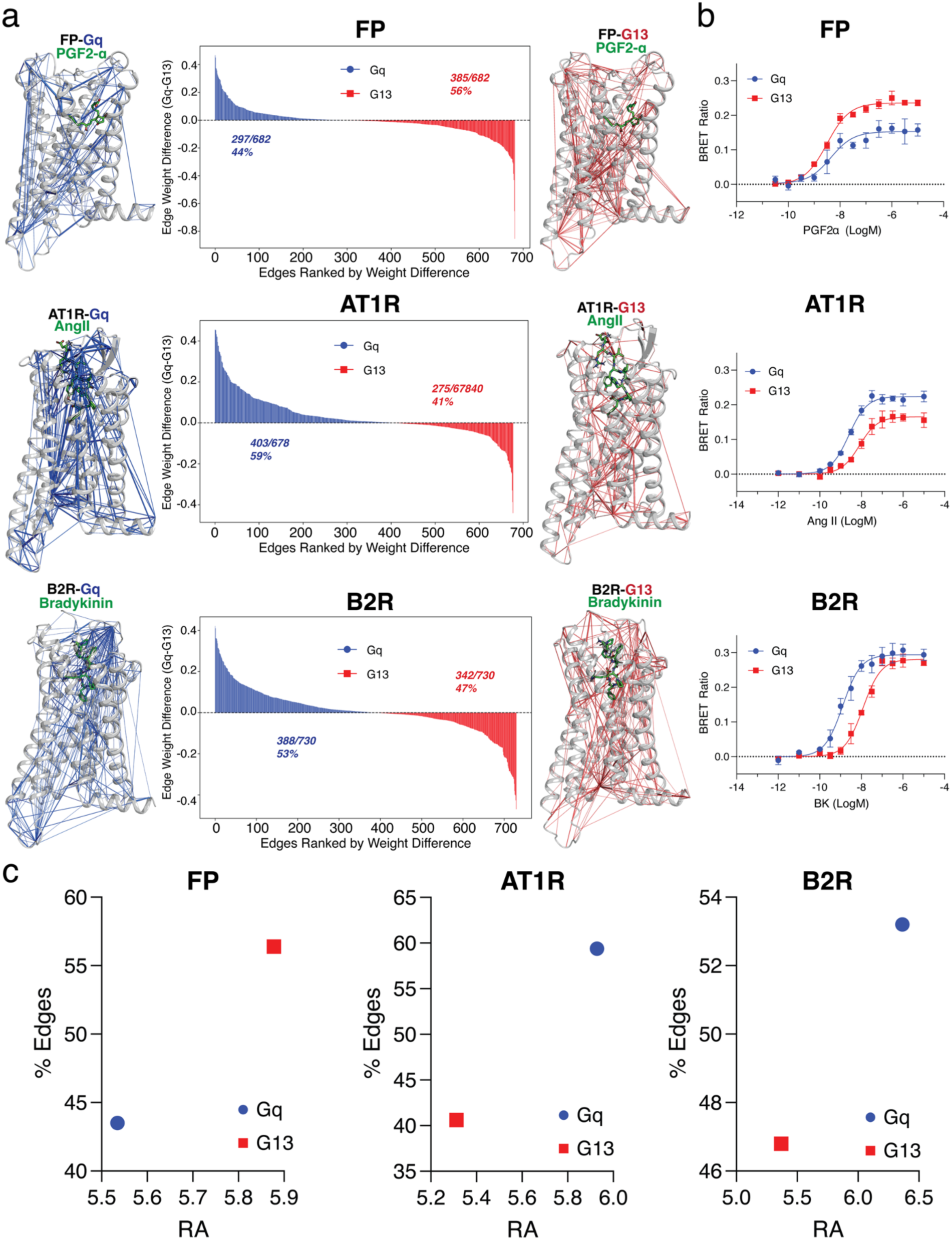
Gα protein-specific differences in Bayesian network edge strengths correlate with coupling preference. **a** Active-state MD-derived Bayesian networks for FP (top), AT1R (middle), and B2R (bottom) in complex with Gα_q_ (blue) or Gα_13_ (red). Edges represent residue-residue dependencies inferred from Bayesian network analysis. Middle panels show the distribution of edge weight differences (Δw = w_Gq_ - w_G13_) ranked across all edges in the universal network for each receptor. Blue bars indicate edges stronger in the Gα_q_-bound state (Δw > 0), and red bars indicate edges stronger in the Gα_13_-bound state (Δw < 0). The percentages denote the fraction of edges stronger in each Gα protein-bound state. **b** Gα protein-specific Rho BRET concentration-response curves for FP (top), AT1R (middle), and B2R (bottom) measured in HEK293TΔGα (lacking endogenous Gα proteins) cells expressing either Gα_q_ (blue) or Gα_13_ (red). Data represent mean ± SEM from three independent experiments. Curves were fit using nonlinear regression. **c** Relative activity plotted against the percentage of edges stronger in each Gα protein-bound state. For each receptor, the Gα protein associated with a larger fraction of stronger edges in the Bayesian network exhibits higher RA.

We next asked whether this network asymmetry parallels functional Gα protein coupling. To test this, FP and AT1R were overexpressed to saturated levels of cell surface expression (Supplementary Fig. 8) in HEK293TΔGα cells, which lack endogenous Gα proteins, allowing us to isolate the maximal capacity of each receptor to activate individual Gα signaling pathways (Gα_q/11_ versus Gα_12/13_). Receptors were co-expressed with either Gα_q_ or Gα_13_ together with the Rho BRET biosensor. This approach enabled measurement of isolated signaling activity for each receptor-Gα pair. The signaling outputs (Fig. 5b) mirrored the differences in allosteric edge strengths observed between the Gα_q_- and Gα_13_-bound states of FP and AT1R (Fig. 5c), and this relationship persisted across varying levels of Gα re-supplementation (Supplementary Fig. 10). To test whether this relationship is generalizable, we extended this analysis to another promiscuous Class A GPCR, the bradykinin B2 receptor (B2R), which also couples to both Gα_q/11_ and Gα_12/13_ like AT1R and FP^6^. MD simulations (Supplementary Fig. 11) followed by BNM analysis on B2R (Supplementary Data 18-20) revealed that, like AT1R, a greater proportion of edges in B2R had stronger co-dependencies in the Gα_q_-bound state than Gα_13_-bound state (Fig. 5a), indicating that the Gα_q/11_ pathway is supported by a more densely interconnected allosteric network than Gα_12/13_ in the B2R. Consistent with the network analysis, the Gα_q/11_ pathway exhibited higher signaling efficiency than the Gα_12/13_ pathway at B2R (Fig. 5b, c; Supplementary Fig. 10). These results quantitatively link Gα protein-specific differences in network edge strengths to coupling preference and hierarchy at promiscuous GPCRs to their cognate G proteins. The Gα protein-bound state enriched in a larger fraction of stronger allosteric edges corresponds to the functionally preferred signaling pathway. Overall, our findings suggest differences in Gα protein-specific allosteric network edge strengths to be the main driver of Gα protein functional selectivity.

## DISCUSSION

How GPCRs encode selectivity and promiscuity toward coupling distinct Gα protein families has remained a central question in the field, particularly given the growing interest in developing biased ligands that target promiscuous GPCRs as therapeutic agents^11–17^. Here, we delineate the structural and dynamic determinants that underlie coupling preferences in two prototypical promiscuous GPCRs, the AT1R and FP receptors, across two major Gα protein classes, Gα_q/11_ and Gα_12/13_. Our findings uncover previously unrecognized ligand- and Gα protein-contact sites unique to either the Gα_q_ or Gα_13_ protein-bound states of FP and AT1R that contribute to selective signaling and show that each receptor engages Gα proteins through uniquely configured allosteric networks across different domains of the receptors. These networks likely impose distinct conformational constraints that shape Gα protein pathway selectivity and confer robustness to mutational perturbations of primary Gα protein coupling, providing mechanistic insight into how selective and biased signaling is encoded at the ligand-receptor-Gα protein level. Importantly, LBS and GPI residues are necessary but not sufficient for Gα protein coupling selectivity. Rather, selectivity is shaped by the embedding of Gα protein-uniquely important residues within broader, receptor-wide allosteric networks. These findings highlight that GPCR allostery links the binding pocket and coupling interface to the entire receptor, not just to each other.

Structural and targeted mutagenesis studies have long established the importance of TMs, the LBS, and the GPI in GPCR-Gα protein activation^1,7,19,42,43^. Our whole receptor alanine-scanning mutagenesis of FP and AT1R, integrated with Gα functional assays and MD simulations, reveals regional sensitivities and previously unrecognized contact sites that govern differential coupling to Gα_q/11_ versus Gα_12/13_. TM6 exerted receptor-specific influences, acting as a leading selectivity determinant in AT1R, consistent with observations in other GPCRs^19^, but not in FP. ECLs were more enriched in Gα protein-uniquely important residues in FP coupling relative to AT1R while ICLs contained enriched clusters of Gα protein-uniquely important residues in both receptors, and the unresolved N-terminal regions displayed opposite regulatory effects, being more important for Gα_q/11_ in FP and Gα_12/13_ in AT1R. The C-termini of GPCRs have been predicted *in silico* to be enriched in signaling features for Gα_q/11_ relative to Gα ^48^. Here, we validated this prediction by experimentally demonstrating that these unresolved structures harbor Gα_12/13_ inhibitory residues at both receptors, supporting a model in which the C-tail restrains Gα_12/13_ coupling relative to Gα_q/11_. Our mutagenesis approach provides unique insight into these disordered regions and their role in Gα protein selectivity, complementing structural studies. It also highlights the critical role of unstructured or disordered regions on receptor function. We have confirmed and further elucidated GPI residues playing an important role in Gα protein selectivity^18,19^. This suggests a complementary mechanism in which a conserved receptor core supports general Gα protein engagement, while distributed networks of residues fine-tune coupling specificity, in conjunction with Gα protein-specific residue contacts within the GPI. We also identified previously unrecognized receptor- and Gα-bound-state-specific ligand contacts within the LBS and GPI of FP and AT1R. These mechanistic insights, revealed only through the integrated analysis of alanine-scanning mutagenesis and MD simulations, can be harnessed to establish a principled framework for the rational design of functionally selective ligands and small molecules targeting the LBS and GPI. Indeed, in an accompanying study^38^, we used these approaches to identify a small, fragment-like negative allosteric modulator of AngII-mediated AT1R activation by predicting key allosteric residues.

To move beyond local interaction network models, we employed BNM to operationally define allostery as probabilistic co-dependencies between spatially distant receptor regions that collectively shape Gα coupling outcomes. BNM revealed receptor-specific architectures of allosteric communication underlying Gα protein coupling. Although BNM infers statistical dependencies rather than direct physical interactions, the predicted network pathways were strongly supported by systematic mutagenesis and functional coupling, both here and previously ^38^, with perturbations at distal network positions producing coupling effects consistent with model predictions. BNM further suggested that strongly weighted, highly connected networks are associated with preferred Gα protein coupling, as observed for AT1R-Gα_q/11_and FP-Gα_12/13_ (Fig. 6). This organization is conceptually analogous to a neural network, in which information processing emerges from the collective activity of multiple interconnected nodes rather than from a single dominant connection. Similarly, a highly interconnected GPCR allosteric network may provide multiple, partially redundant routes linking the LBS to the GPI, thereby reducing dependence on individual residues and maintaining productive ligand-receptor-G protein communication despite local perturbations. In contrast, weaker or sparser networks rely on fewer critical nodes and communication routes, making secondary coupling modes, such as AT1R- Gα_12/13_ and FP-Gα_q/11_, more susceptible to disruption by individual mutations. Thus, the architecture of ligand-receptor-G protein-specific allosteric networks may explain both preferential coupling and differential mutational resilience, with robust coupling emerging from multiple interconnected routes rather than from a single dominant allosteric pathway. Beyond providing this mechanistic framework, BNM enables the identification and hierarchical ranking of residues contributing to these networks, thereby revealing candidate allosteric sites through which Gα protein coupling may be selectively modulated. The extension of this approach to B2R further supports its broader applicability across GPCRs. Within this framework, AT1R and B2R preferentially couple to the Gα_q/11_ family, consistent with previous classifications^49–51^, whereas FP preferentially couples to Gα_12/13_, refining its G protein coupling profile.

**Figure 6.**
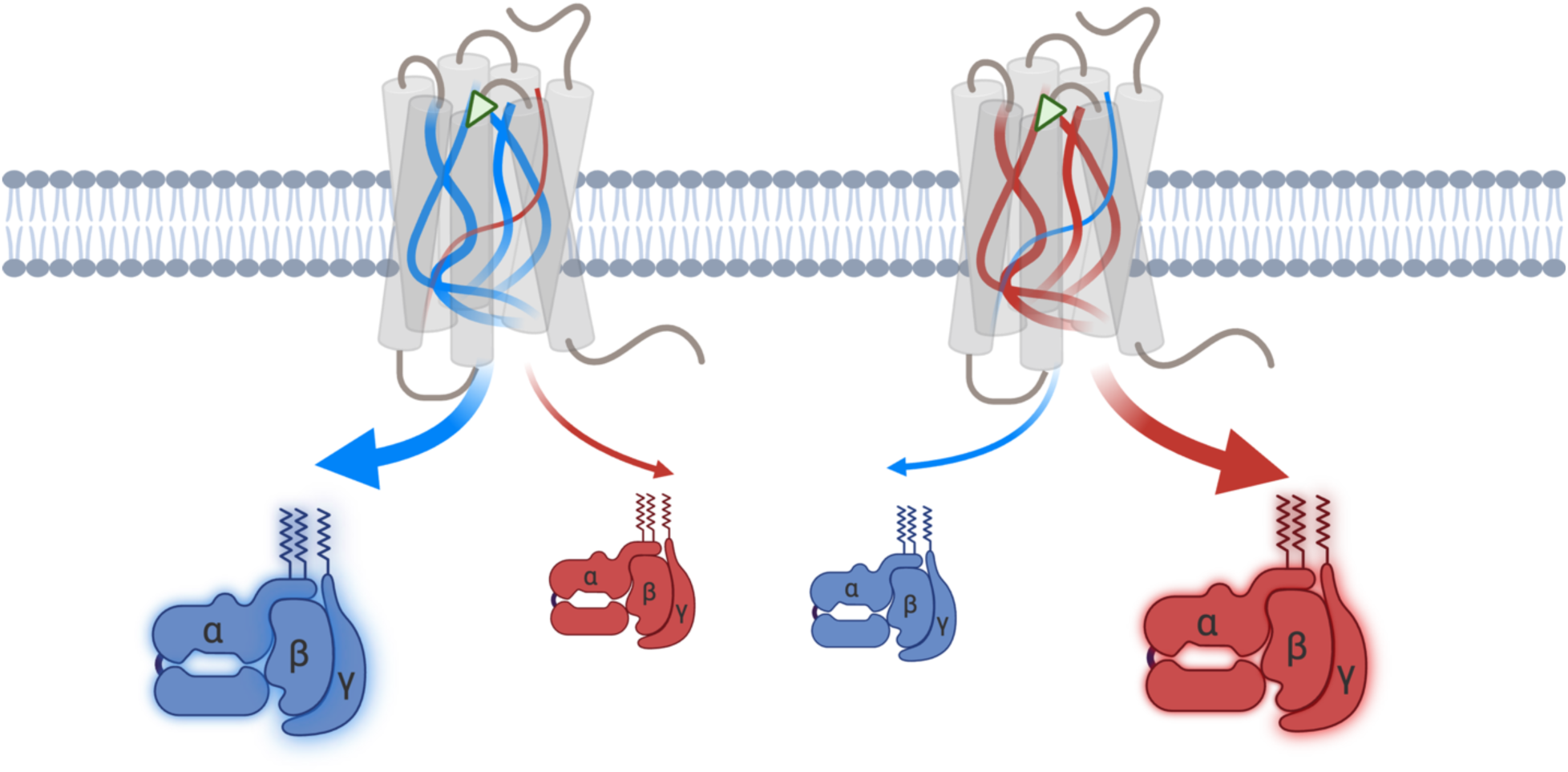
Gα protein coupling selectivity at promiscuous GPCRs is associated with allosteric network density and efficiency. Schematic representation of predominantly Gα_q/11_ -coupled (left) and Gα_12/13_-coupled (right) promiscuous GPCRs. The predominantly Gαq/11-coupled receptor exhibits greater density and efficiency of its Gα_q/11_ (blue) allosteric network relative to its Gα_12/13_ (red) network, resulting in more robust Gα_q/11_ coupling, whereas the inverse is observed for the predominantly Gα_12/13_-coupled receptor.

Several additional considerations and potential caveats warrant discussion. Our functional analyses were performed using alanine or glycine substitutions in the context of the full-length receptor; substitutions with alternative residues may yield distinct functional outcomes. Moreover, residues that did not display penetrant effects on Gα protein coupling in these assays may nonetheless contribute to the propagation of allosteric information across the residue network, despite having limited standalone effects. Importantly, mutations that preserved overall receptor expression and general coupling capacity frequently altered Gα protein preference, arguing against expression level or assay amplification as primary drivers of the observed selectivity patterns. In our BNM analysis, changes in network connectivity between inactive and active receptor-Gα protein coupling states were inferred from comparisons of receptor-Gα protein complexes versus unbound receptor states and did not explicitly account for Gα protein nucleotide state (GDP versus GTP), which may differentially shape allosteric communication. Our findings complement rather than replace structure-based models, suggesting that static GPCR-G protein interfaces define permissive coupling, while dynamic allosteric organization determines coupling preference and robustness. Finally, while our hierarchical ranking of receptor-Gα protein coupling preferences was assessed using a common downstream effector of Gα_q/11_ and Gα_12/13_, namely Rho, coupling hierarchies may differ when evaluated using alternative downstream effectors.

In summary, by integrating probabilistic network inference with functional data from an ensemble of receptor mutants, we establish a mechanistic framework in which Gα protein selectivity at promiscuous GPCRs emerges from receptor-specific hierarchies of intra-receptor communication within a shared signaling architecture. In this framework, we revise the prevailing view of FP as a predominantly Gα_q/11_ coupled receptor, showing instead that its functional coupling is primarily Gα_12/13_, reflecting differences in Gα protein-specific allosteric networks. We further identify the residue-level determinants underlying this Gα protein functional selectivity, providing the molecular basis for understanding, and manipulating, Gα protein coupling selectivity through strategies such as biased ligand design or receptor mutagenesis. More broadly, our validated *in silico* approaches, provides a mean to resolve coupling hierarchies in promiscuous GPCRs. Overall, this work reframes Gα protein selectivity as an emergent property of receptor-wide allosteric network architecture, rather than a simple consequence of local interface compatibility.

## METHODS

### Reagents

Dulbecco’s Modified Eagle Media (DMEM), fetal bovine serum (FBS), and other cell culture additives were purchased from Gibco, Life Technologies (Carlsbad, CA). Linear polyethylenimine MW 25000 (PEI) was purchased from Polyscience, Inc (Warrington, PA). PGF2α was purchased from Cayman Chemical (Ann Arbor, MI). Coelenterazine 400a was purchased from Nanolight Technology (Pinetop, AZ). Anti-HA-peroxidase rat antibody (3F10) was purchased from Roche (Manheim, Germany). BSA was purchased from Fisher BioReagents (Hampton, NH). Chemiluminescence reagents were purchased from Perkin-Elmer (Waltham, MA). SIGMAFAST OPD, 16% paraformaldehyde (PFA), and pertussis toxin (PTX) were purchased from Sigma-Aldrich (St. Louis, MO). Q5 high-fidelity DNA polymerase, Gibson Assembly Mastermix, DpnI, and other PCR reagents were purchased from New England BioLabs (Ipswich, MA). Clean-up and 96-well miniprep kits were purchased from Bio Basic (Markham, ON).

### Cloning of FP alanine scan library

A two-part PCR strategy was employed for whole receptor mutagenesis, as described previously^26,52,53^. Briefly, site-directed mutagenesis primers were generated with 18 bp of Gibson of homology for Gibson assembly recombination to insert alanine/glycine mutations in FP and were ordered from Integrated DNA Technologies (Coralville, IA). The mutations were introduced through a step-down PCR; two separate PCRs were run to split the vector in half. Each half of the PCR sample was combined and digested with DpnI. Purified samples containing the two half vectors were then assembled by Gibson assembly. The re-annealed vector was transformed into bacterial colonies, and one colony was picked and amplified. DNA vectors were harvested using a 96 well miniprep kit and sent for sequencing to verify successful mutagenesis (Genome Québec).

### DNA constructs

Origin of replication primers were described previously^26,52,53^. WT and mutant HA-tagged FP receptors containing a signal peptide were constructed in pcDNA3.1. Combined mutations were generated using pre-existing mutants as templates, utilizing the same PCR strategy mentioned above. PKC, Rho, Gα_q_ Gαβγ and Gα_13_ Gαβγ BRET biosensors, as well as pcDNA3.1 were described previously^34,44^.

### Cell culture and transfections

HEK293T, HEK293SL, Gα_q/11_ knock-out (KO)^54^, Gα_12/13_ KO^55^, and Gα-null^56^ cells were cultured in DMEM supplemented with 10% FBS, and 20 μg/ml gentamicin. Cells were authenticated (GeneCopoeia™) and tested for mycoplasma contamination periodically (ABM™, Mycoplasma PCR Detection kit). Cells were grown at 37°C in 5% CO_2_ and 90% humidity. Cells were seeded at a density of 2.0×10^4^ cells per well in a white 96-well flat bottom plate (for BRET) or clear 96 well flat bottom plate (for ELISA) and transiently transfected with receptor and sensor DNA using PEI transfection reagent. Briefly, a total of 1 μg of DNA in 100 μl of PBS was mixed with 3 μl of 1 mg/ml PEI in 100 μl of PBS. For FP Gα_q/11_ signaling, 150 ng of FP DNA with 20 ng of PKN-RlucII DNA and 80 ng of rGFP-CAAX DNA were used in HEK293TΔGα_12/13_ cells or 150 ng of FP DNA, 30 ng of Gα_q_-RlucII, 120 ng of Gβ_1_, and 120 ng of Gγ_1_-GFP10 were used in HEK293T cells. For FP Gα_12/13_ signaling, 150 ng of FP DNA with 20 ng of PKN-RlucII DNA and 80 ng of rGFP-CAAX DNA were used in Gα_q/11_ KO cells or 30 ng of FP DNA, 48 ng of Gα_13_-RlucII, 120 ng of Gβ_1_, and 120 ng of Gγ_1_-GFP10 were used in HEK293T cells. For AT1R Gα_q/11_ signaling, 60 ng of AT1R DNA with 60 ng of PKC-c1b sensor DNA were used in HEK293SL cells or 150 ng of AT1R DNA with 20 ng of PKN-RlucII DNA and 80 ng of rGFP-CAAX DNA were used in Gα_12/13_ KO cells. For AT1R Gα_12/13_ signaling, 150 ng of AT1R DNA with 20 ng of PKN-RlucII DNA and 80 ng of rGFP-CAAX DNA were used in HEK293SL cells in the presence of 500 nM YM-254890 (Gα_q/11_ inhibitor) or in Gα_q/11_ KO cells. For isolated Rho signaling of GPCR-Gα protein pairs, 150 ng of FP DNA, AT1R DNA, or B2R DNA was co-transfected with 2.5, 5, 7.5, 10, or 12.5 ng of either Gα_q_ or Gα_13_ as well as 20 ng of PKN-RlucII DNA and 80 ng of rGFP-CAAX DNA in Gα-null cells. To assess the contribution of Gα_i/o_ to FP and AT1R Rho signaling, the respective signaling assays were performed under identical conditions in the absence or presence of pertussis toxin (PTX). For ELISA, 150 ng of HA-FP DNA and 150 ng of spFLAG-AT1R DNA, respectively, were used in HEK293T cells and HEK293SL cells, respectively, and increasing amounts (50-250 ng) of HA-FP DNA, HA-AT1R DNA, and B2R DNA. Empty pcDNA was used to make up 1 µg of the total DNA amount. After 20-min incubation, the DNA/PEI complexes were dispensed into cells in 96-well plates (15 µl/well). Twenty-four hours later, the medium was replaced and all assays were performed 48 h post-transfection.

### BRET measurements

Cells were seeded at a density of 2.0×10^4^ cells per well in polyornithine coated white 96 well flat bottom plate and transiently transfected with receptor and sensor DNA. 48 hours post-transfection, cells were incubated for 1 h with Tyrode’s buffer (140 mM NaCl, 2.7 mM KCl, 1 mM CaCl_2_, 12 mM NaHCO_3_, 5.6 mM D-glucose, 0.5 mM MgCl_2_, 0.37 mM NaH_2_PO_4_, 25 mM HEPES, pH 7.4). Cells were stimulated for 2 min 30 s with serially diluted of PGF2α from 10^-11^ M to 10^-5^ M and AngII and BK from 10^-12^ M to 10^-5^ M and the signal was recorded using the Biotek Synergy 2 plate reader with filter sets of 410/80 nm (donor) and 515/30 nm (acceptor). Cell-permeable substrate coelenterazine 400a (final concentration of 2.5 μM) was added 3 min prior to BRET measurements BRET ratios were calculated by dividing acceptor emission by donor emission. The data was fitted to 12-point concentration-response curves and analyzed for its activity.

### Cell-surface ELISA

Clear 96-well flat bottom plates were coated with poly-L-lysine. WT and mutant receptors were transfected into HEK293SL cells. On the day of the experiment, the media was removed and the cells were washed with PBS and fixed with 4% PFA. The cells were then blocked with 2% BSA and incubated with anti-HA HRP (1:1000). The cells were washed and 100 μl SIGMAFAST OPD solution was added to each well. After 10 min, 25 μl 3 M HCl was added to stop the reaction. Absorbance at 492 nm was measured using a BioTek Synergy 2 plate reader.

### Experimental data analysis and statistics

Statistical analyses were performed using GraphPad Prism 11. Twelve-point BRET concentration-response data from each replicate were fitted to the Hill equation using three-parameter nonlinear regression, with the Hill slope fixed at 1. Within each replicate, raw BRET responses were normalized to the span of the WT response, with 0% and 100% defined by the bottom and top best-fit values of the WT curve, respectively. For AT1R, data were transformed and analyzed as previously described^26^. Following normalization and transformation, concentration-response curves were fitted by nonlinear regression to derive best-fit values for logEC_50_, span (Emax), and log(Emax/EC_50_), defined as relative activity (RA). For concentration-response curves with very low response amplitudes (Emax <10% of WT), logEC_50_ could not be reliably estimated because of the limited dynamic range of the response. Accordingly, logEC_50_ and RA values were considered undetermined and excluded from subsequent analyses for mutants with Emax <10%. For replicates in which nonlinear regression did not yield a stable curve fit, Emax was estimated from the response at the highest agonist concentration tested (10^-5^ M), whereas logEC_50_ and RA were considered undetermined. For comparisons between WT and mutant pharmacological parameters, unpaired two-tailed t-tests were performed. To account for multiple comparisons, *p*-values were converted to *q*-values using the adaptive two-stage step-up procedure of Benjamini, Krieger and Yekutieli^57^, controlling the false discovery rate (FDR) at *Q* = 5%.

### MD Simulations

The following 3D structures were used as starting structures for MD simulation in this study: fully active state of AT1R-Sar1-AngII-Gα_q_-Gβ_1_-Gγ_2_ complex (Protein Data Bank (PDB) accession number: 7F6G), inactive state of AT1R-ZD7 complex (PDB ID: 4YAY), active state of FP receptor-PGF2α-Gα_i/s/q_(chimera)-Gβ_1_-Gγ_2_-scFv16-Nb35 complex (PDB ID: 8IUK), active state of B2R-bradykinin-Gα_q_-Gβ_1_-Gγ_2_ complex (PDB ID: 7F6H). The structure of GPR35-Lodoxamide-Gα_13_-Gβ_1_-Gγ_2_-scFv16 structure (PDB ID: 8H8J) was used to model the Gα_13_ the FP and AT1R complexes Protein preparation and minimization of all WT models based on the above-described 3D experimental structures were performed in Schrödinger Maestro^58^. The AT1R-AngII-Gα_q_-Gβ_1_-Gγ_2_ model was created by swapping the residues in Sar1-AngII to wildtype AngII. The soluble cytochrome b562 was removed in the inactive state structure of AT1R, three AT1R mutations were reverted to wildtype. A 10-residue gap including ICL3, and two 4-residue gaps in ECL2 were modeled onto the structure using the Homology Modeling module in Schrödinger Maestro^61^. The FP receptor-PGF2α-Gα_q_-Gβ_1_-Gγ_2_ model was created by removing the scFv16 and Nb35 and swapping the residues of Gαi/s/q(chimera) to WT Gα_q_. The mutations in the Gαq of 7F6H were reverted to the WT sequence (Uniprot sequence P50148). The antagonist-bound inactive FP receptor was generated by integrating the Br-derivative (8VL, from PDB ID: 5YHL) structure with the inactive-state model retrieved from GPCRdb^59^. All models were created using Homology Modeling module in Schrödinger Maestro^58^. For generating the Gα_13_ bound structures we overlaid the Gα_13_-Gβ_1_-Gγ_2_ bound AT1R, FP receptor and B2R models, AT1R-AngII, FP receptor-PGF2α, and B2R-bradykinin were superimposed with the GPR35-Lodoxamide-Gα_13_-Gβ_1_-Gγ_2_-scFv16 complex after which we removed the GPR35-Lodoxamide-scFv16 from the complex in PyMOL^60^. This resulted in AT1R-AngII-Gα_13_-Gβ_1_-Gγ_2_, FP receptor-PGF2α-Gɑ_13_-Gβ_1_-Gγ_2_, and B2R-bradykinin-Gα_13_-Gβ_1_-Gγ_2_ models respectively. The remaining mutations in the models were reverted to wildtype, and missing domains/loops were recreated using the Prime module^61^ within Schrödinger Maestro^61^ with the exception of the ɑ-helical domain in FP receptor-Gα_q_ and FP receptor-Gα_13_ and AT1R-Gα_13_ models. Next, the missing sidechains and hydrogen atoms were added, and histidine protonation states were assigned. Finally, the models were energetically minimized using the standard settings in the protein preparation wizard module^61^. We generated our MD simulation systems using GROMACS^62^ package (version 2022) with the Chemistry HARvard Molecular Mechanics (CHARMM36) force field for proteins^63^. Receptor complexes were positioned in a membrane bilayer through the orientation of proteins in membranes (OPM) database. Next, a pre-equilibrated POPC (palmitoyl-oleoyl-phosphatidylcholine) bilayer, ions, and CHARMM Transferable Intermolecular Potential with 3 Points (TIP3P) water^64^ as solvent were generated around the complex to generate each simulation box. The final system dimensions were as follows: AngII-AT1R-Gα_q_-Gβ_1_-Gγ_2_ 120 × 120 × 179 Å, including 371 lipids, 58170 waters and 150 mM NaCl; AngII-AT1R-Gα_13_-Gβ_1_-Gγ_2_ 120 × 120 × 167 Å, including 375 lipids, 53186 waters and 150 mM NaCl; ZD7-AT1R 75 × 75 × 122 Å, including 130 lipids, 13,801waters and 150 mM NaCl; PG2α-FP-Gα_q_-Gβ_1_-Gγ_2_ 120 × 120 × 165 Å, including 381 lipids, 52472 waters and 150 mM NaCl; PG2α-FP-Gα_13_-Gβ_1_-Gγ_2_ 120 × 120 × 163 Å, including 383 lipids, 51767 waters and 150 mM NaCl; 8VL-FP 90 × 90 × 114 Å, including 199 lipids, 18027 waters and 150mM NaCl; BK-B2R-Gα_q_-Gβ_1_-Gγ_2_ 150 × 150 × 157 Å, including 612 lipids, 77801 waters and 150 mM NaCl; BK-B2R-Gα_13_-Gβ_1_-Gγ_2_ 130 × 130 × 167 Å, including 453 lipids, 62741 waters and 150 mM NaCl. The simulation systems, once prepared, underwent an initial minimization with position restraints of 10 kcal/mol·Å^2^ on all heavy atoms encompassing the protein, ligand, and lipids. This was succeeded by a 1-ns heating phase that escalated the temperature from 0 K to 310 K under the NVT ensemble, utilizing the Nosé-Hoover thermostat. Subsequently, the system was subjected to an equilibration simulation within the NPT ensemble. Here, the initial 1 ns had the 10 kcal/mol·Å^2^ position restraint. This restraint was then progressively reduced: first to 5 kcal/mol·Å^2^ and then diminishing to 1 kcal/mol·Å2 in decrements of 1 kcal/mol·Å^2^. Each decrement was accompanied by a simulation spanning 5 ns. The final step of the equilibration involved a 100 ns unrestrained NPT ensemble. Next, a total of 5 production MD simulation (each with a different random starting velocity), each 1 μs long was carried out, with the simulation snapshots stored every 20 ps. To assess the convergence of MD simulations the root-mean-square deviation (RMSD) of the TM backbone was calculated using GROMACS rms using the following AT1R residues (Supplementary Fig. 4, 9): TM1: 25-57, TM2: 62-90, TM3: 98-131, TM4: 142-166, TM5: 190-229, TM6: 235-268 and TM7: 274-305; FP receptor residues: TM1: 30-61, TM2: 65-94, TM3: 106-138, TM4: 149-173, TM5: 196-233, TM6: 243-275 and TM7: 283-307; B2R residues: TM1: 55-85, TM2: 90-120, TM3: 126-161, TM4: 170-196, TM5: 217-257, TM6: 262-299, TM7:303-335. All the simulation frames were used to calculate MD convergence. True positive rates for ligand-residue and Gα protein-residue contacts were calculated by cross-referencing identified contacts from simulations and experimentally affected residues from our alanine scans.

### MD contact analysis and definition of LBS and GPI residues

Intermolecular contacts were analyzed independently for five replicate MD trajectories of the AT1R-Gα_q_, AT1R-Gα_13_, FP-Gα_q_, and FP-Gα_13_ complexes. Dynamic contacts were identified using GetContacts (https://getcontacts.github.io/) with the corresponding topology and trajectory files. All contact types implemented by GetContacts were considered using the --*itypes all* option, and identical contact definitions and geometric criteria were applied to all systems. Separate calculations were performed for contacts between the receptor and agonist and between the receptor and Gα subunit. Multiple atom-atom contacts, or contacts with multiple partner residues within the same frame, were treated as a single occupied frame for that receptor residue and were therefore not counted additively. Frequencies ranged from 0, indicating that no contact was observed, to 1, indicating that a contact was present throughout the analyzed trajectory.

Contact frequencies were first calculated separately for each of the five replicate trajectories. The mean frequency was then calculated as the arithmetic mean of the five run-specific frequencies. Because equal numbers of frames were analyzed from each run, this mean was equivalent to the frequency obtained from the concatenated trajectory; the values were cross-checked against the corresponding GetContacts frequency output from the concatenated trajectory.

Ligand-binding-site (LBS) residues were defined as receptor residues that contacted the agonist-AngII for AT1R or PGF2α for FP- and satisfied either of the following criteria: (i) a mean contact frequency of ≥ 0.20 across the five runs or (ii) a contact frequency of ≥ 0.10 in at least four of the five runs. The second criterion retained lower-occupancy contacts that were reproducibly observed across independent simulations. G-protein-interface (GPI) residues were defined using the same criteria based on contacts between receptor residues and the corresponding Gα_q_ or Gα_13_ subunit. For receptor-level comparisons and structural representations, the LBS and GPI sets were defined as the union of qualifying residues identified in the Gα_q_- and Gα_13_-coupled simulations for each receptor.

All receptor residues for which a contact was detected were retained in the per-run heatmaps for completeness. Residues that failed both the mean frequency and replicate-reproducibility criteria were indicated by red residue labels and were not included in the final classified LBS or GPI sets. Contact frequency heatmaps and residue-level comparisons were generated using in-house Python scripts, and structural representations were prepared in PyMOL^60^.

### Calculation of residue interaction energies

The nonbonded interaction energy (Eij) between any two residues (i and j) is composed of two separate energy terms: The van der Waals (vdW) interaction energy (VLJ) given by the Lennard-Jones (LJ) potential (Eq. 1) and the electrostatic interaction energy (VC) given by Coulombic potential (Eq. 2). Specifically, the short-range (within 12 Å) Coulombic and van der Waals forces were calculated using *gmx energy* module in GROMACS^59^ and extracted from the energy log file. The computed interaction energy is defined as the sum of the LJ and Coulombic interaction energies (Eq. 3) for every MD snapshot (25,000). All favorable interaction energies are <0 kJ/mol and hence have a negative sign.

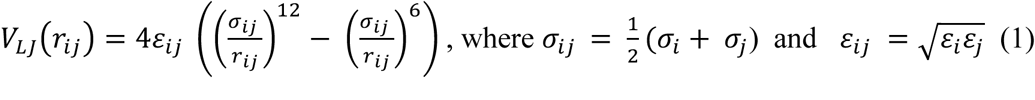

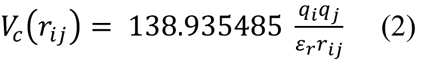

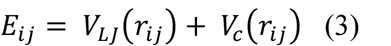

### Construction of differential Bayesian network models

DBNM was used as a proof-of-concept approach to compare two systems by constructing separate Bayesian network (BN) models for each condition (i.e., active vs. inactive AT1R or FP). For each network, we computed the weighted degree of node i as *D_i_* = ∑_j∈Г_*_i_*w_ij_, where Γ_i_is the set of neighbors of the node i and *w_i_*_j_ is the commensurate, universal-scale, edge strength^37^. Prior work has established weighted degree as a robust proxy for functional importance, with highly connected nodes often corresponding to residues critical for protein function or stability^20^. To identify differential network features, we defined the weighted degree difference between systems A and B as Δ*D_i_*(G_A_, G_B_) = *D_i_*(*A*) − *D_i_*(*B*) the weighted degree difference for the node i between the two BNs G_A_, G_B_. This measure quantifies the extent of changes in the dependency structure as identified by BNs in the two systems, directly pointing to the most important differential modulator(s) for these two systems (illustrated in Chen *et al*.^38^).

### Construction of individual and universal Gα_q/11_ vs. Gα_12/_ Bayesian network models (BNMs)

The residue-wise interaction energy data from Gα_q/11_ and Gα_12/13_ ensemble trajectories were concatenated for AT1R, FP, and B2R, respectively, and used for generating the universal Gα_q/11__Gα_12/13_ BNM graph for each receptor using the BaNDyT software^45^. Independent Gα_q/11__Gα_12/13_ BNM models were generated for each receptor using a search-and-score approach.

The Minimum Uncertainty (MU) scoring function was employed, with 100 random restarts to ensure convergence^37^. Then the weight for each edge in Gα_q/11_ and Gα_12/13_ systems was respectively re-computed taking the universal network topology, but using mutual information calculated from Gα_q/11_ dataset only or Gα_12/13_ only. This yielded different Gα_q/11_ and Gα_12/13_ graphs with the same graph topology, but different weights across all edges. This analysis was used to identify salient co-dependencies that differentiate between the two Gα_q/11_-bound vs. Gα_12/13_-bound states. To that effect, we define Δ*W_i_*(G_G**α**q/11_, G_G**α**12/13_) = *W_i_*(Gαq/11) − *W_i_*(Gα12/13) to system-specific importance of each edge.

### Enrichment factor calculation

Residues were ranked by their differential weighted degree (ΔWD) values as Δ*Wd_i_*(G_active_, G_inactive_) = *Wd_i_*(active) − *Wd_i_*(inactive), calculated as the difference between the weighted degree in the active-state BN model and the inactive-state BN model. Experimental “positives” were defined as residues showing a statistically significant change (p < 0.05, Student’s t-test) in at least one pharmacological parameter (EC_50_, Emax, or relative activity) compared to wild-type in the alanine scanning mutagenesis assays. For each integer n from 1 to the total number of ranked residues, the enrichment score (ES) was computed as: 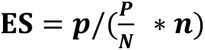, where ***n*** is sample size of the subset; ***N*** is total sample size; ***p*** is true positive rate of the subset; ***P*** is total true positive rate. An ES > 1 indicates that the top-n residues are enriched for experimental positives relative to the background distribution. Enrichment curves were plotted by sequentially calculating ES for increasing n values, with every sixth point shown for visualization (Supplementary Fig. 6).

### Fisher’s Exact Test for Enrichment Analysis

To assess statistical significance of enrichment at specific ranked subsets, Fisher’s exact test was performed comparing the number of experimental positives and negatives in the subset versus the remainder of the ranked list. The 2 × 2 contingency table consisted of: (i) Experimental positives in subset (ii) Experimental negatives in subset (iii) Experimental positives in remainder (iv) Experimental negatives in remainder. A one-tailed Fisher’s exact test (p < 0.05) was used to determine whether the proportion of positives in the subset was greater than expected by chance.

## DATA AVAILABILITY

The molecular MD simulation snapshots for all the simulations in this work are deposited in the GPCRMD database. All other data generated and analyzed are included in this article or its Supplementary Information.

## CODE AVAILABILITY

BaNDyT is available on GitHub at: https://github.com/bandyt-group/bandyt. Or upon reasonable request.

## ACKNOWLEDGEMENTS

We acknowledge our discussions with Dr. Ning Ma, Dr. Supriyo Bhattacharya, and Dr. Babgen Manookian. Funding for this work was supported by NIH grants R35 GM156498 to N.V., and R01 LM013876 to N.V., A.S.R and S.B. and Canadian Institutes of Health Research grants PJT-162368 and PJT-173504 to S.A.L.

## AUTHOR CONTRIBUTION

T.S.B., H.C., A.C, W.J.C.v.d.V., Y.N., Y.C., N.V., and S.A.L conceptualized the study. T.S.B., H.C., A.C., W.J.C.v.d.V., Y.N., Y.C., F.M.H., A.S.R., S.B., N.V., and S.A.L. established the methodology of the study. A.C. and F.C., T.S.B generated and characterized the FP and AT1R alanine mutants signaling and performed data analysis. H.C. and W.J.C.v.d.V performed MD simulations and BNMs of receptors. J.Y.P performed data analysis. T.S.B., H.C., J.Y.P and W.J.C.v.d.V performed data interpretation. T.S.B. and H.C. prepared the manuscript. N.V. and S.A.L. revised and edited the final version of the manuscript. N.V. and S.A.L. acquired funding and supervised the study.

