## Supplementary Figures 1-11 for "Allosteric pathways govern Gα protein coupling selectivity at promiscuous GPCRs"

### TITLE

[1]Department of Medicine, Research Institute of the McGill University Health Centre, McGill University, Montréal, QC, Canada. [2]Department of Computational and Quantitative Medicine, Beckman Research Institute of the City of Hope, Duarte, CA, USA. [3]Irell and Manella Graduate School of Biological Sciences, Beckman Research Institute of the City of Hope, Duarte, CA, USA. [4]Department of Pharmacology and Therapeutics, McGill University; Montréal, QC, Canada. [5]Department of Biochemistry and Molecular Medicine, Institute for Research in Immunology and Cancer, Université de Montréal, Montreal, QC, Canada.

These authors contributed equally to this work: Tegvir S. Boora, Hanyu Chen

These authors jointly supervised this work: Nagarajan Vaidehi, Stéphane A. Laporte

*

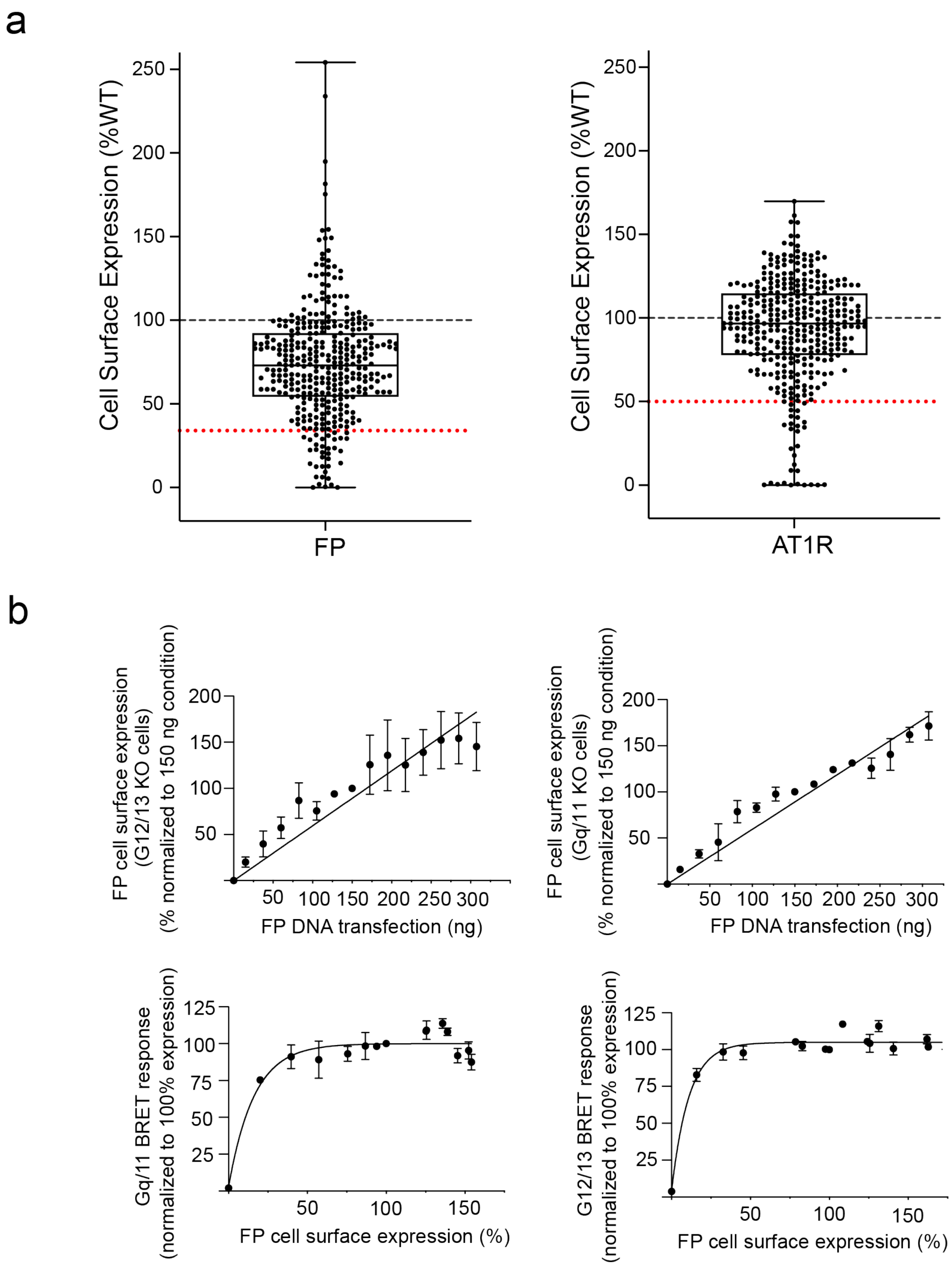


**Supplementary Figure 1. Cell surface expression of FP and AT1R mutants and titration of wild-type (WT) FP DNA transfection amount against cell surface receptor expression and maximal BRET response. a** Cell surface expression of all 358 FP mutants (left) was assessed using ELISA. Previously reported cell surface expressions of the AT1R mutant library using ELISA are shown on the right. **b** Increasing amounts of FP DNA were transfected into HEK293 cells to quantify receptor expression by ELISA, under conditions matched to those used for BRET measurements of $G\alpha_{q/11}$ coupling (left) in $G\alpha_{12/13}$ KO cells and $G\alpha_{12/13}$ coupling (right) in $G\alpha_{q/11}$ KO cells. Expression levels were normalized to 100% at 150 ng of transfected DNA. **c** Maximal BRET responses for (left) $G\alpha_{q/11}$ and (right) $G\alpha_{12/13}$ were measured following stimulation with 1 μM $PGF_{2}\alpha$ and plotted as a function of the corresponding FP receptor expression levels determined in panels **a** and **b**, respectively. Data represent the mean ± S.E.M. of triplicates from three independent experiments.

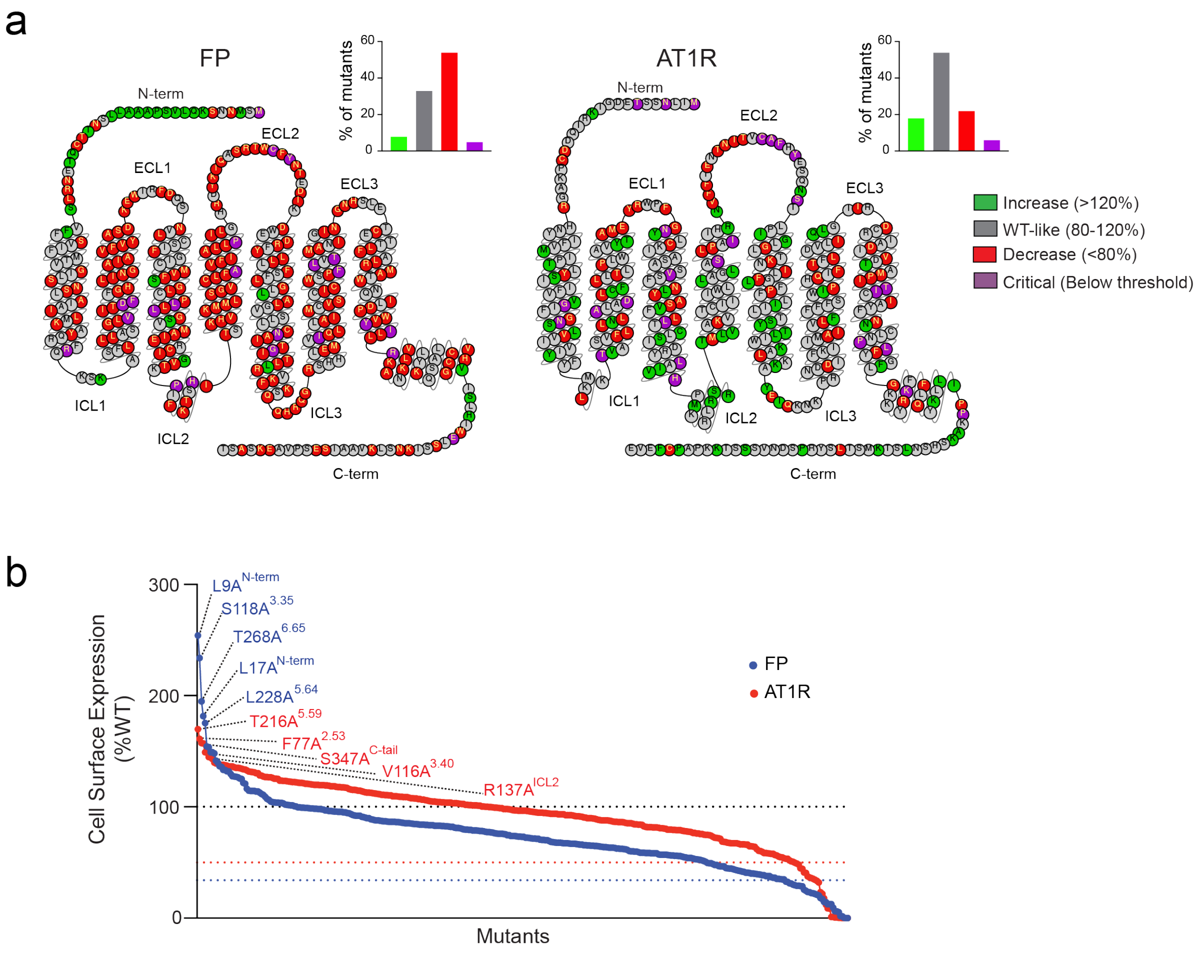


**Supplementary Figure 2. Domain-specific regulation of receptor cell surface expression. a** Effects of alanine substitution on cell surface expression of FP (left) and AT1R (right). Snakeplots highlight mutations that increased (green) expression relative to WT receptor (120%), maintained WT-like expression (grey), decreased relative expression but above critical threshold (<80%), and mutations that severely decreased relative cell surface expression below the critical expression threshold for signaling established for each receptor (34% for FP and 50% for AT1R). **b** Waterfall plot of the effects of receptor mutations on cell-surface expression. expression, with WT expression set to 100% (black dotted line). The five mutations with the highest expression are highlighted for each receptor. Colored dotted lines indicate the receptor expression thresholds required for sufficient signaling (34% of WT for FP, blue; 50% of WT for AT1R, red).

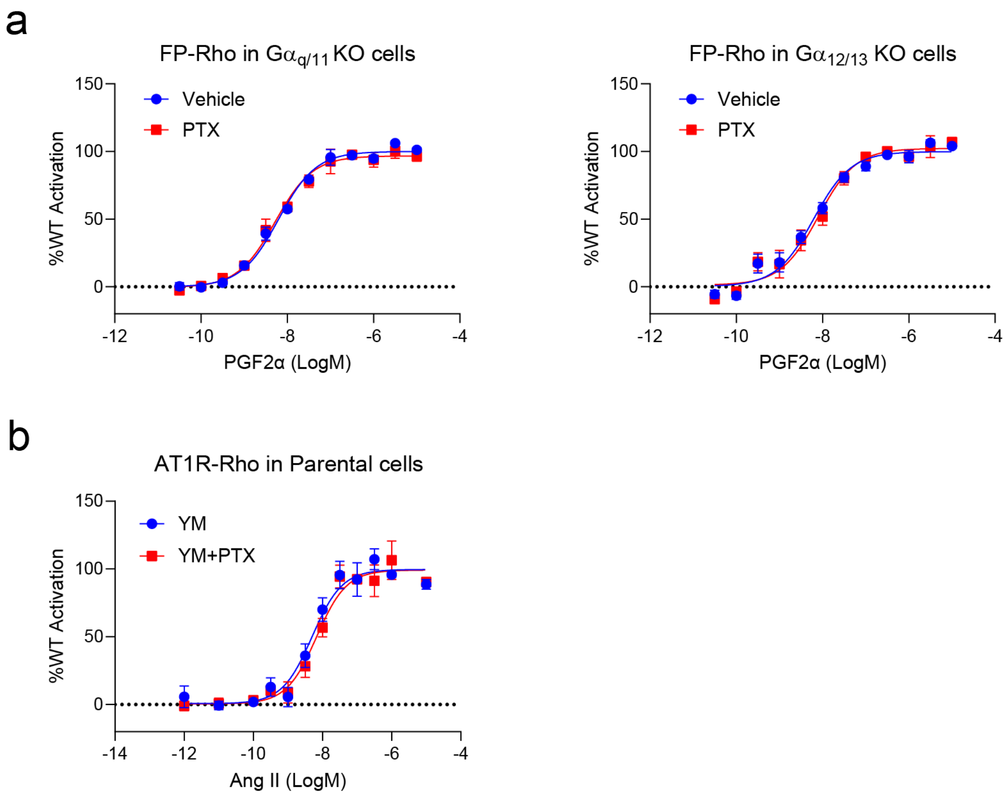


**Supplementary Figure 3. Extent of Gi/o involvement in Rho signaling as assessed with BRET-based biosensor. a** FP and Rho biosensor transfected G$\alpha_{q/11}$ KO cells and G$\alpha_{12/13}$ KO cells were preincubated with either vehicle or pertussis toxin (PTX, Gi/o inhibitor) for ~18 hr then stimulated with increasing concentrations of PGF2α. **b** AT1R and Rho biosensor transfected parental HEK293T cells were stimulated with increasing concentrations of AngII along with YM (Gq/11 inhibitor) in the absence or presence of preincubation with PTX. Data are presented as mean ± SEM from three independent experiments, each performed in triplicate. Curves were generated using GraphPad Prism.

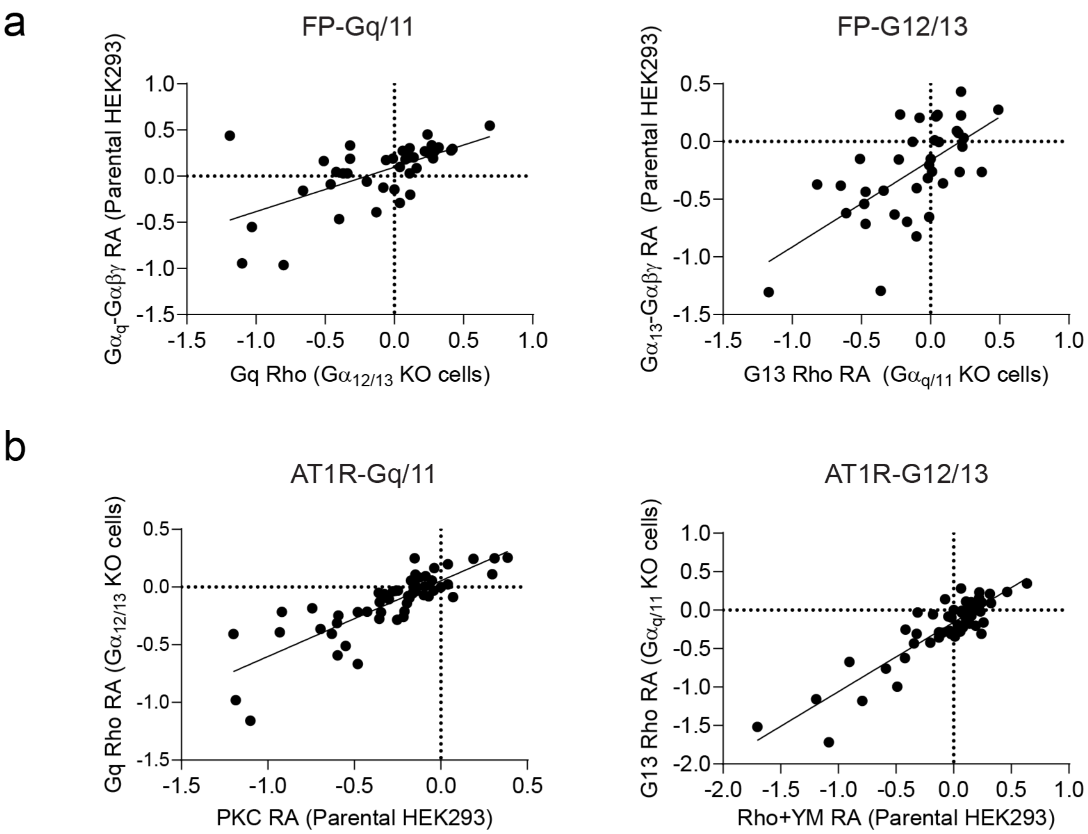


**Supplementary Figure 4 Cross-validation of Gα protein coupling measured using distinct BRET-based biosensors. a** RA (Δlog(Emax/$EC_{50}$) values were calculated from the Emax and $EC_{50}$ values of the $G\alpha_q$ Gαβγ biosensor in HEK 293 cells and Rho biosensor in $G\alpha_{12/13}$ KO cells and between $G\alpha_{13}$ Gαβγ BRET biosensor (left) and Rho BRET in $G_{q/11}$ KO cells of WT and 41 select FP alanine mutants (right). Data are represented as the mean ± S.E.M. of three independent experiments. (A) Slopes of 0.4809 ± 0.1014 and $R^2$ value of 0.3719 (left) and of 0.1589 ± 0.2524 and $R^2$ value of 0.3728 (right) were obtained from a simple linear regression. The $G\alpha_q$ and $G\alpha_{13}$ Gαβγ responses recapitulate the responses of Rho sensor in $G\alpha_{12/13}$ KO cells and $G\alpha_{q/11}$ KO cells. **b** RA (Δlog(Emax/$EC_{50}$) values were calculated from the Emax and $EC_{50}$ values of the PKC in HEK 293 cells and Rho sensor in $G\alpha_{12/13}$ KO cells and between Rho BRET biosensor in presence of $G\alpha_{q/11}$ inhibitor in HEK 293 cells and Rho BRET in $G\alpha_{q/11}$ KO cells of WT and AT1R 54 select alanine mutants. Data are represented as the mean ± S.E.M. of three independent experiments. Slopes of 0.68 ± 0.06 and $R^2$ value of 0.6791 (left) and of 0.7990 ± 0.06 and $R^2$ value of 0.7990 (right) were obtained from a simple linear regression. The PKC and Rho sensors in HEK 293 cells recapitulate the responses of Rho sensor in $G\alpha_{12/13}$ KO cells and $G\alpha_{q/11}$ KO cells, respectively.

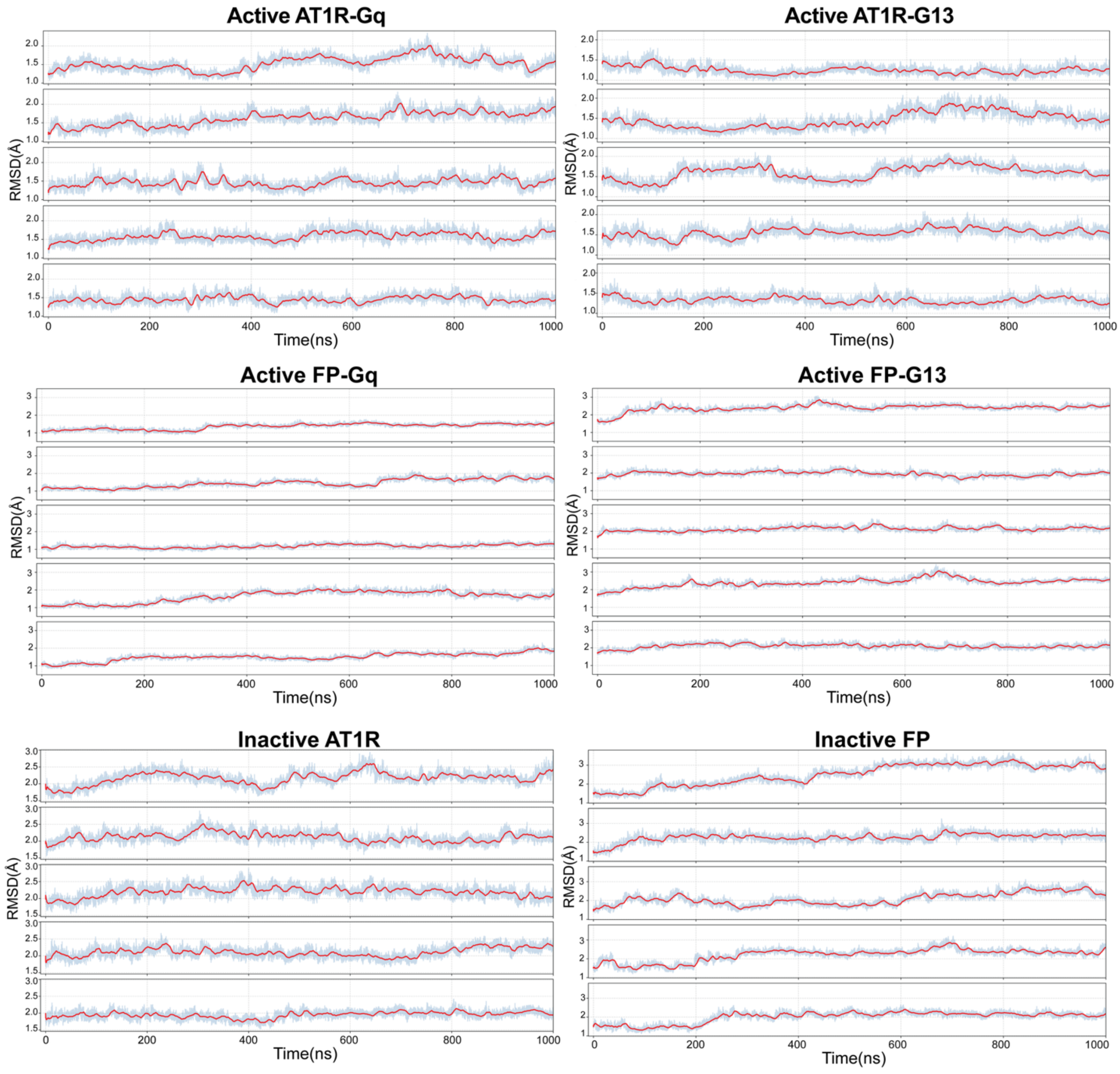


**Supplementary Figure 5. RMSD analysis of MD trajectories.** Root-mean-square deviation (RMSD) of transmembrane backbone Cα atoms is shown for five independent 1-μs MD simulation trajectories for each system. Active-state complexes of AT1R and FP bound to $G\alpha_q$ or $G\alpha_{13}$ are shown, along with inactive-state simulations. For each trajectory, the light blue trace represents the instantaneous RMSD relative to the starting structure, and the red line indicates the running average. RMSD values plateau over time in all systems, indicating structural stability of the simulated complexes throughout the production runs.

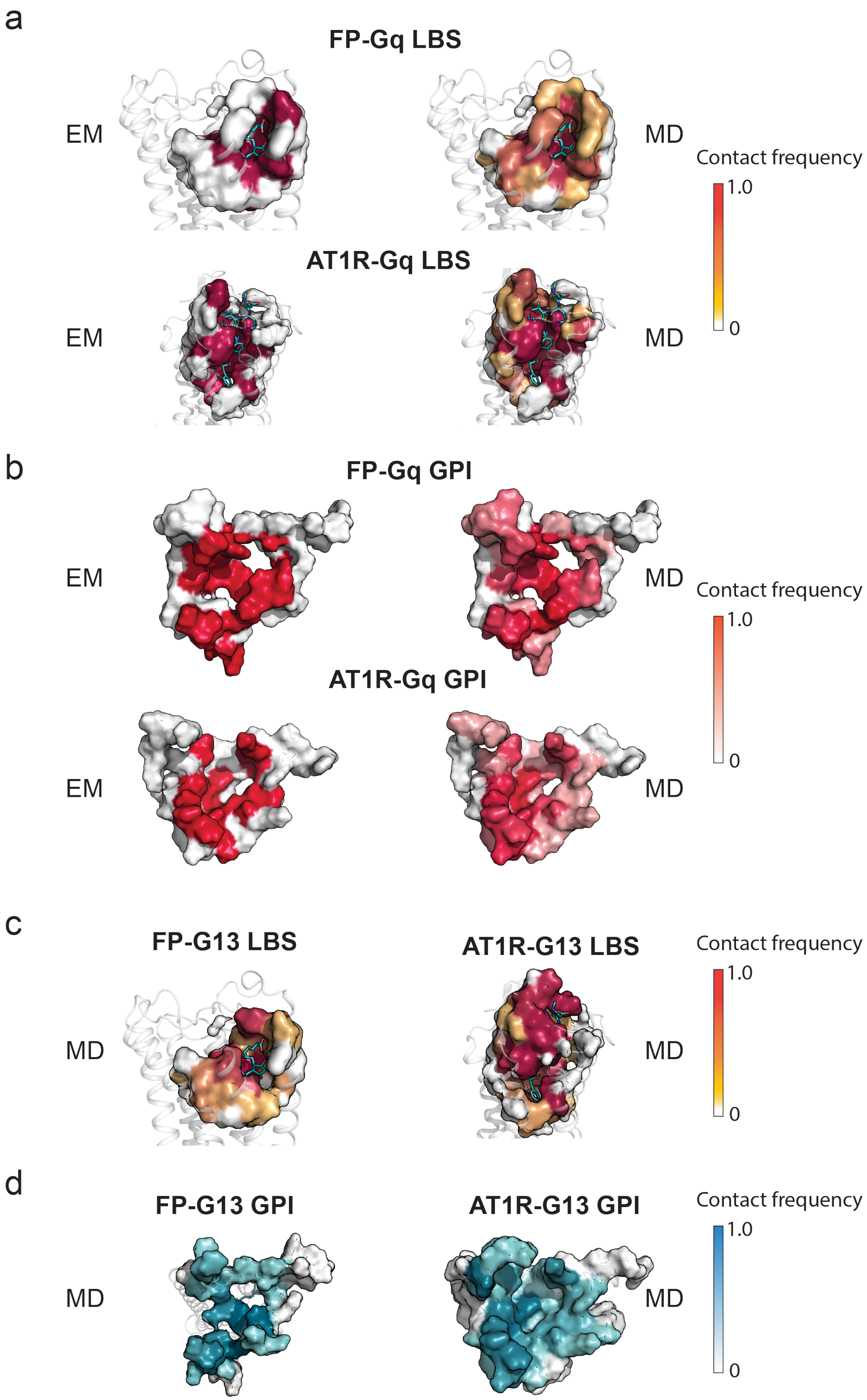

a
FP-Gq LBS
EM
MD
Contact frequency
1.0
0
AT1R-Gq LBS
EM
MD
b
FP-Gq GPI
EM
MD
Contact frequency
1.0
0
AT1R-Gq GPI
EM
MD
c
FP-G13 LBS
AT1R-G13 LBS
Contact frequency
1.0
0
MD
MD
d
FP-G13 GPI
AT1R-G13 GPI
Contact frequency
1.0
0
MD
MD

**Supplementary Figure 6. Molecular dynamics (MD) simulations uncover previously unknown low frequency residue-contacts with the ligand binding sites (LBS) and G protein-binding interface (GPI) of FP and AT1R. a** MD simulations recapitulate ligand-residue contacts identified in the $G\alpha_q$-bound and ligand-bound crystal structures (left) of FP (top) and AT1R (bottom) as high frequency contact but further uncover lower frequency contacts (right). **b** MD simulations recapitulate Gα protein-residue contacts identified in the $G\alpha_q$-bound crystal structures (left) of FP (top) and AT1R (bottom) as high frequency contact but further uncover lower frequency contacts (right). **c** MD simulations uncover ligand-residue contacts for the $G\alpha_{13}$ and ligand-bound states of FP (left) and AT1R (right). **d** MD simulations uncover Gα protein-residue contacts for the $G\alpha_{13}$-bound state of FP (left) and AT1R (right).

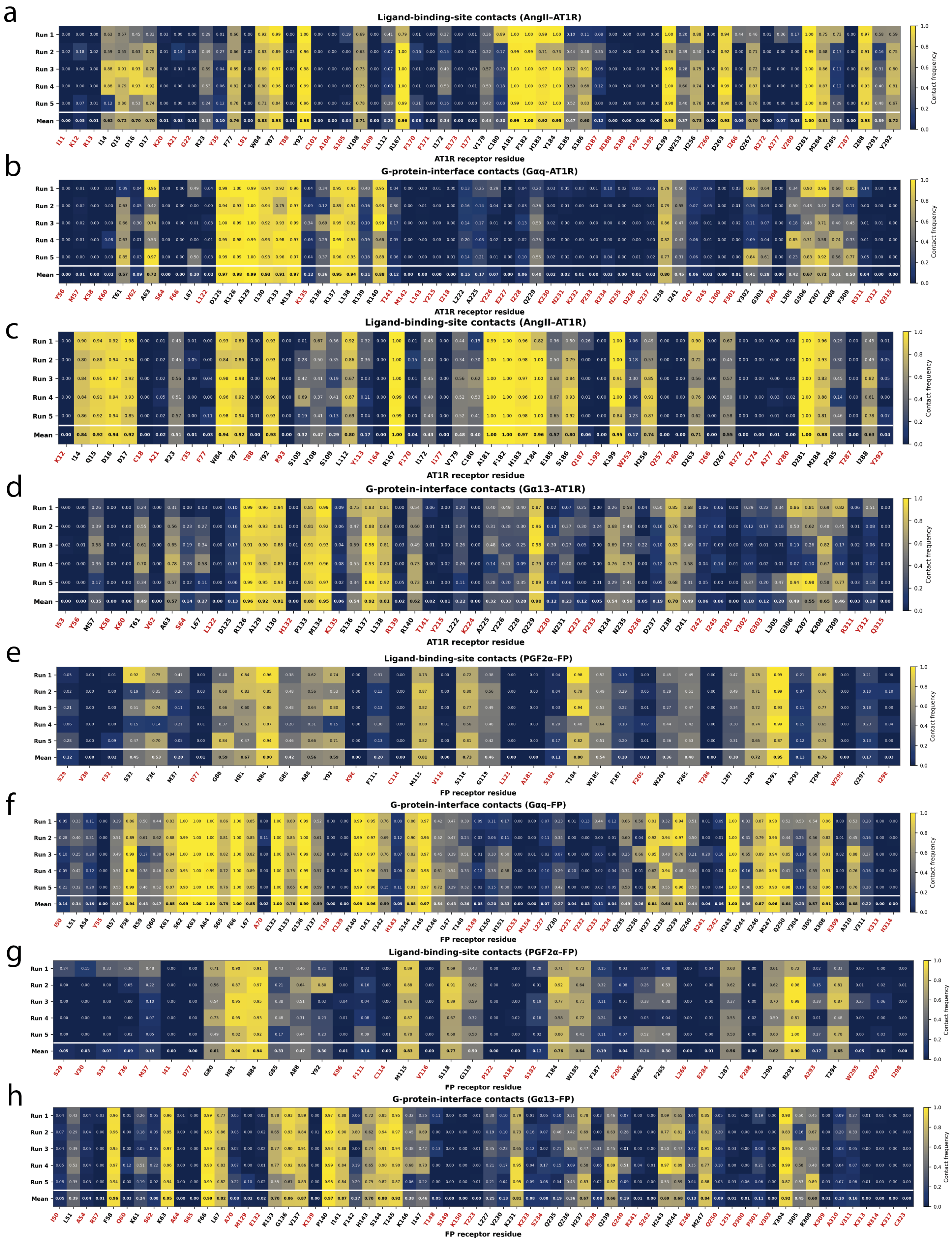


**Supplementary Figure 7. Per-run contact-frequency heatmaps for ligand-binding-site and G-protein-interface residues in FP and AT1R complexes coupled to Gα$_q$ or Gα$_{13}$. a** AngII-

AT1R ligand-binding-site (LBS) contact frequencies in the AT1R-G$\alpha_q$ complex. **b** G$\alpha_q$-AT1R G-protein-interface (GPI) contact frequencies. **c** AngII-AT1R LBS contact frequencies in the AT1R-G$\alpha_{13}$ complex. **d** G$\alpha_{13}$-AT1R GPI contact frequencies. **e** PGF2α-FP LBS contact frequencies in the FP-G$\alpha_q$ complex. **f** G$\alpha_q$-FP GPI contact frequencies. **g** PGF2α-FP LBS contact frequencies in the FP-G$\alpha_{13}$ complex. **h** G$\alpha_{13}$-FP GPI contact frequencies. Each heatmap shows contact frequencies from five independent MD runs and their means. Values range from 0 (no contact) to 1 (contact throughout the trajectory). Residue labels shown in red did not meet the contact-frequency and reproducibility criteria defined in Methods; these residues are displayed for completeness.

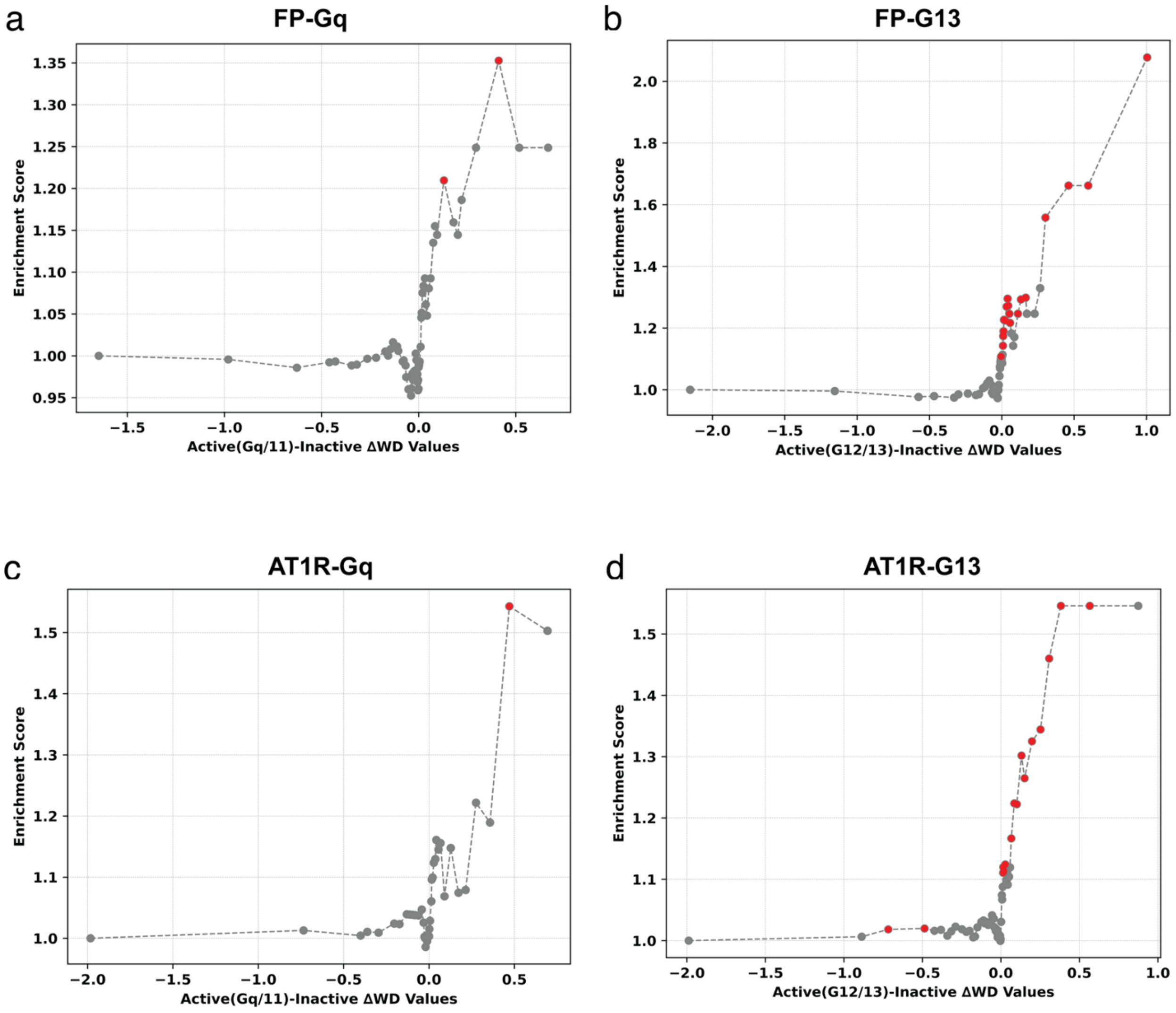


**Supplementary Figure 8. Enrichment analysis of $G\alpha_{q/11}$-/$G\alpha_{12/13}$- and inactive-state Bayesian Network (BN) predictions across residue-ranked thresholds.** Enrichment curve of residues ranked by ΔWD values (every sixth dot shown). Each dot represents the enrichment score at a given *n* (top *n* residues ranked by ΔWD), comparing the proportion of true positives within the subset to the expected background rate. Red dots correspond to subset sizes that reach statistical significance (Fisher's exact test $p < 0.05$) (see Methods for more details). The enrichment curve shown are respectively ranked by **a** $\Delta Wd_i(\mathcal{G}_{\text{FP\_Gq/11}}, \mathcal{G}_{\text{FP\_inactive}})$; **b** $\Delta Wd_i(\mathcal{G}_{\text{FP\_G12/13}}, \mathcal{G}_{\text{FP\_inactive}})$; **c** $\Delta Wd_i(\mathcal{G}_{\text{AT1R\_Gq/11}}, \mathcal{G}_{\text{AT1R\_inactive}})$; **d** $\Delta Wd_i(\mathcal{G}_{\text{AT1R\_G12/13}}, \mathcal{G}_{\text{AT1R\_inactive}})$ cross-referencing respective alanine/glycine scanning experimental positives.

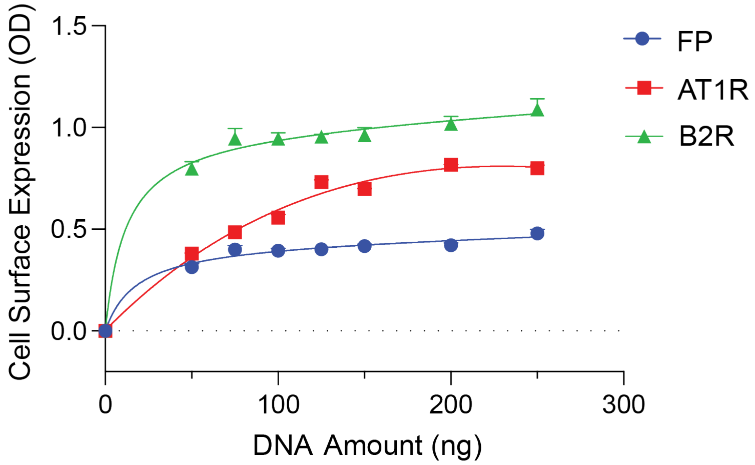


**Supplementary Figure 9. Relationship between transfected receptor DNA and cell-surface expression.** Increasing amounts of FP, AT1R, or B2R DNA were transfected into cells, and cell-surface receptor expression was quantified by ELISA. Data are presented as mean ± SEM from three independent experiments, each performed in triplicate.

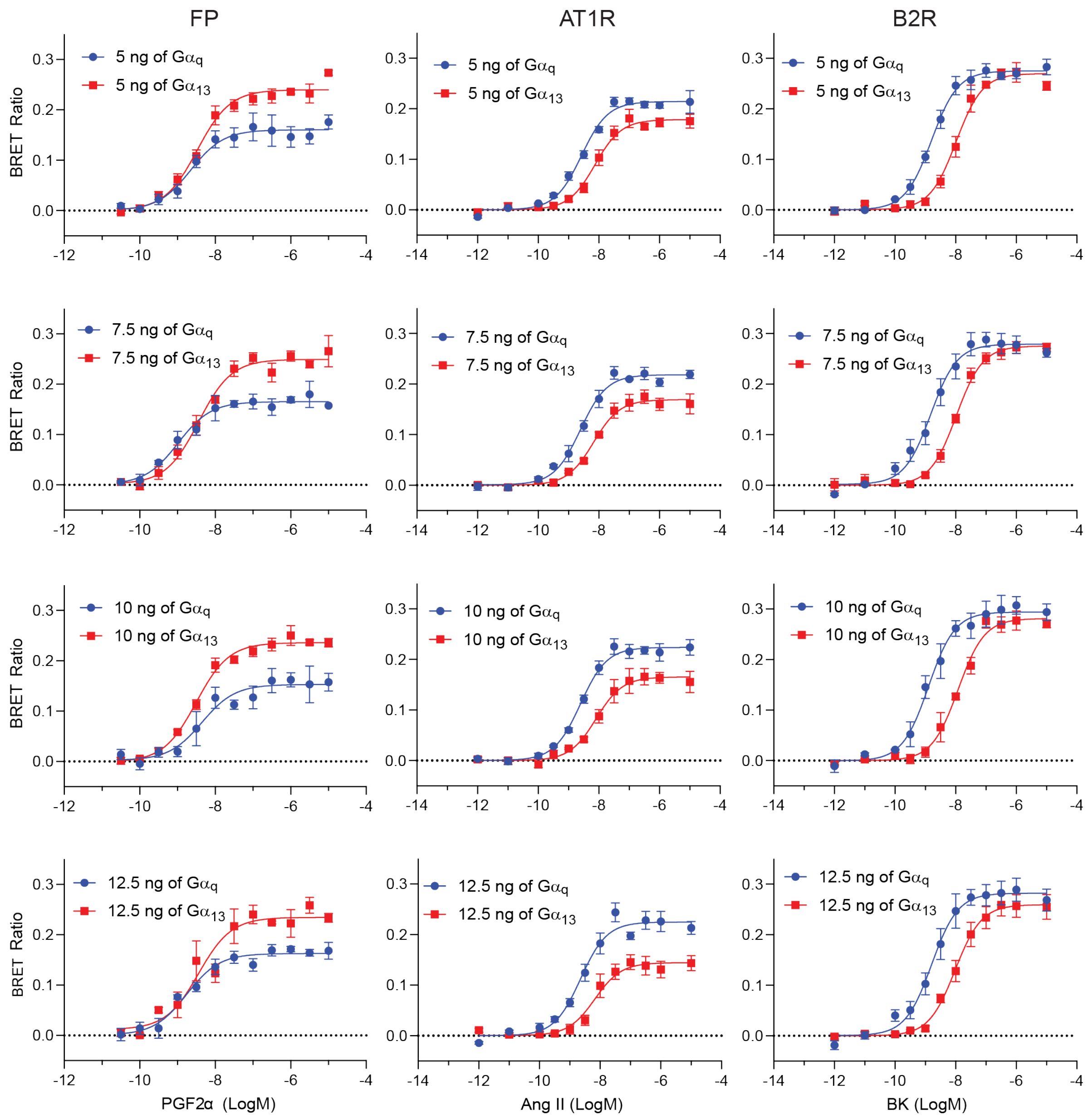


**Supplementary Figure 10**. **Receptor–Gα protein signaling hierarchy is maintained across varying Gα protein expression levels**. Increasing amounts of $G\alpha_q$ and $G\alpha_{13}$, respectively, were co-transfected with receptor DNA (FP, AT1R, or B2R) and Rho BRET biosensor in Gα-null cells which were stimulated by respective endogenous ligand (PGF2α, Ang II, BK) to obtain 12 dose point-response curves for FP (left), AT1R (middle) and B2R (right). Curves plotted using GraphPad Prism. Data represent the mean ± S.E.M. of triplicates from three independent experiments.

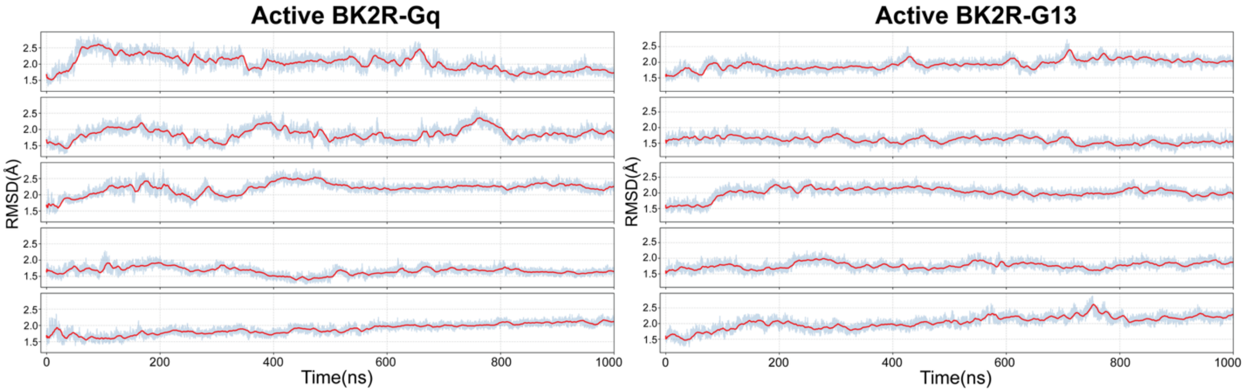


**Supplementary Figure 11. RMSD analysis of MD trajectories.** Root-mean-square deviation (RMSD) of transmembrane backbone Cα atoms is shown for five independent 1-μs MD simulation trajectories for each system. Active-state complexes of B2R bound to $G\alpha_q$ or $G\alpha_{13}$ are shown. For each trajectory, the light blue trace represents the instantaneous RMSD relative to the starting structure, and the red line indicates the running average. RMSD values plateau over time in all systems, indicating structural stability of the simulated complexes throughout the production runs.
